# *Plasmodium berghei* is resistant to aryl amino acetamides that inhibit *P. falciparum* growth by targeting the phospholipid transfer protein *Pf*START1

**DOI:** 10.64898/2026.09.22.753445

**Authors:** Claudia B.G. Barnes, Mrittika Chowdury, Dawson B. Ling, Thomas McLean, Alysha H. Literski, Natalie A. Counihan, Oliver Looker, Thorey K. Jonsdottir, Coralie Boulet, William Nguyen, Madeline G. Dans, Emma Yuxin Mao, Alan F. Cowman, Niall D. Geoghegan, Kelly L. Rogers, Danny W. Wilson, Brendan S. Crabb, Stephen W. Scally, Brad E. Sleebs, Hayley E. Bullen, Tania F. de Koning-Ward, Paul R. Gilson

## Abstract

In a previous screen for compounds that inhibit *Plasmodium falciparum* merozoite invasion of red blood cells, we identified the Medicines for Malaria Venture compound MMV006833. This compound inhibits *Pf*START1, a protein implicated in the expansion of the nascent parasitophorous vacuole membrane following invasion, to accommodate the developing ring-stage parasite. Live-cell lattice light-sheet microscopy of invading merozoites revealed that mNeonGreen-tagged *Pf*START1 is released from structures within the merozoite into the nascent parasitophorous vacuole approximately 109 seconds after invasion. Expansion microscopy of *Pf*START1-HA merozoites further showed that these punctate *Pf*START1-containing structures do not colocalise with known secretory organelles (rhoptries, micronemes and dense granules). Although analogues of MMV006833 are highly potent against *P. falciparum*, they were previously found to be ineffective against *P. berghei* parasites in the mouse malaria model. Here, we demonstrate that *Pb*START1 is highly resistant to MMV006833 and its analogues when expressed in *P. falciparum*, indicating that structural differences between the orthologous proteins reduce inhibitor potency. The crystal structure of *Pf*START1 in complex with WEHI-991 revealed the molecular basis for inhibition and provided a structural explanation for the reduced potency of this family of compounds against *P. berghei*. To sensitise *P. berghei* parasites to MMV006833 analogues, the parasites were engineered to express *Pf*START1; however, these chimeric parasites remained insensitive to the compounds. This suggests that factors beyond target engagement, such as compound half-life or bioavailability, contribute to the lack of efficacy observed in the mouse malaria model.

## Introduction

Despite major reductions in morbidity and mortality over the past two decades, malaria remains a devastating global health problem that in 2024 was responsible for 282 million clinical cases and 610,000 deaths^1^. The majority of these were attributable to *Plasmodium falciparum* parasites. The use of antimalarial drugs remains the only option to eliminate parasite infections, with artemisinin combination therapies (ACTs) being the most widely used medications. Concerningly, parasite resistance to artemisinin and its derivatives, as well as partner drugs (e.g. lumefantrine, piperaquine) is spreading rapidly and threatening to reverse recent progress towards malaria elimination^1^. Genetic mutations associated with resistance to ACTs have been detected in many countries, including several in Africa where the impact of malaria is the greatest^2–6^. We therefore urgently need to discover and develop new antimalarial drugs to replace ACTs as their efficacy declines.

*Plasmodium* parasites grow and reproduce inside human red blood cells (RBCs). The parasite’s asexual reproductive cell cycle involves the formation of invasive merozoite-stage parasites that exit the host cell and invade new RBCs. As the invasive merozoite propels itself inside the RBC, it is enveloped in the RBC plasma membrane, which transforms into the parasitophorous vacuole membrane (PVM) after invasion^7^. Soon after invasion, the merozoite expands into a larger, amoeboid ring-stage parasite with the PVM expanding to accommodate it.

One emergent drug target of interest is the *P. falciparum* STeroidogenic Acute Regulatory protein-related lipid Transfer domain-containing protein PF3D7_0104200 (also called *Pf*PV6)^8^. *Plasmodium* parasites have four START domain-containing proteins, of which PF3D7_0104200 (referred to herein as START1) is the most extensively studied. *Pf*START1 appears to be required for expansion of the newly invaded parasite after it enters human RBCs^8–11^ and conditional deletion of *Pfstart1* arrests ring-stage development, indicating that *Pf*START1 is essential^8,12^. *Pf*START1 is a phospholipid transfer protein which is thought to shield the hydrophobic lipid molecules within a central cavity and chaperone their transport between membranes^13^. Several compounds, such as the aryl amino acetamide WEHI-991, have been developed that target *Pf*START1 and inhibit the growth of cultured *P. falciparum* parasites with single-digit nanomolar EC_50_s^10,14^. *Pf*START1 inhibitors likely act by competing for the lipid binding pocket of the protein, thereby preventing phospholipid transport.

Both the conditional deletion of the *Pfstart1* gene using the DiCre recombinase system and its inhibition with WEHI-991 render newly invaded merozoites incapable of expanding into amoeboid ring-stage parasites, thus preventing subsequent development and eventually causing parasite death^8,10,12,15^. *Pf*START1 appears to be expressed throughout the asexual cell cycle, but its localisation is only well-studied in young rings where it concentrates in the parasitophorous vacuole (PV) at the tips of the ring’s amoeboid arms^11^. As *Pf*START1 localises to the PV, it is thought to play a crucial role in facilitating PVM expansion by shuttling lipids from the merozoite to the PVM shortly after RBC invasion^8,10,11^. It is not known if merozoites secrete *Pf*START1 into the PV during or after invasion and so here, we have tagged the protein near its N-terminus with mNeonGreen (NG) and observed the fusion protein following merozoite invasion of RBCs and throughout the subsequent cell cycle.

As clinical antimalarials should ideally target multiple parasite stages^16^, we previously investigated whether WEHI-991 affected parasite invasion of liver cells and mosquito tissues in addition to RBCs^10^. Although WEHI-991 did not inhibit *P. falciparum* gamete development, it inhibited ookinete formation within the mosquito gut, indicating potential as a transmission blocking drug^10^. WEHI-991 did not, however, inhibit the invasion of *P. berghei* sporozoites into cultured hepatocytes^10^. We also analysed the ability of the WEHI-991 analogue WJM-715 to clear or substantially reduce the parasitaemia of *P. berghei*-infected mice following oral administration^14^. WJM-715 was administered for four consecutive days at 50 mg/kg but failed to suppress the parasitaemia of *P. berghei*-infected mice^14^.

Two possible scenarios could account for the failure of WJM-715 to reduce parasite growth in mice. First, the compound’s metabolic liabilities, which were flagged in *in vitro* assays with liver microsomes^10,14^, may have resulted in its rapid elimination *in vivo*. The second possibility is that the structure of *Pb*START1 is sufficiently different from *Pf*START1 that the *Pf*START1 inhibitors do not work efficiently against *P. berghei*. To understand the molecular basis for compound selectivity against *P. berghei* parasites (and *P. falciparum* parasites with mutations in *Pf*START1), we obtained an X-ray structure of *Pf*START1 in complex with WEHI-991 that revealed the amino acid differences between *Pf*START1 and *Pb*START1 underpinning the reduced binding of aryl acetamides to *Pb*START1.

To further explore this in a biological context, we report on replacing the START domain in *P. falciparum* parasites with its *P. berghei* counterpart. These *Pf/Pb*START1 parasites were highly resistant to WEHI-991 and all other analogues tested, indicating this might be why the WEHI-991 analogue WJM-715 failed to limit the growth of *P. berghei* parasites in mice. Therefore, with the aim of sensitising *P. berghei* parasites to the WEHI-991 analogue WJM-715, we replaced the START domain of the *P. berghei* orthologue with the *P. falciparum* START domain and evaluated the compound’s efficacy in mice infected with these chimeric *Pb/Pf*START1 parasites. These chimeric parasites were resistant to the compound, suggesting that pharmacokinetic factors, as well as structural differences between *Pf*START1 and *Pb*START1, explain the lack of efficacy in mice.

## Results

### *Pf*START1 is found in different locations throughout the asexual blood stage

*Pf*START1 possesses a PEXEL motif which has been shown to be cleaved by the ER-resident plasmepsin V (PMV) protease^8^. For most PEXEL proteins, this facilitates export across the PVM of the intraerythrocytic parasite and into the RBC compartment, via the *Plasmodium* translocon of exported proteins (PTEX)^17–19^. However, despite being a substrate of PMV, is has been shown that *Pf*START1 is retained in the PV rather than being exported into the RBC cytoplasm^8^. Similarly, cell fractionation and protease protection experiments have demonstrated that *Pf*START1 resides in the parasite and the PV at the schizont stage^10^. *Pf*START1’s inability to be exported is likely attributable to the presence of a charged N-terminal lysine residue on the protein at the P_1_’ position after PMV cleavage which presumably inhibits its interaction with PTEX^8^.

*Pf*START1 has recently been tagged upstream of its START domain with mNeonGreen (NG) and was shown to localise to the tips of the amoeboid arms in early ring-stage parasites^11^; however, the localisation of *Pf*START1 throughout the asexual blood stage has not been systematically investigated. Therefore, we N-terminally appended NG to *PfS*TART1 by inserting NG’s coding sequence downstream of a 5’ *Pfstart1* homology block containing the RILKE PEXEL motif (Fig 1A). This was fused to the rest of the *Pfstart1* gene that had been recodonised to prevent recombination in this region. A T2A skip peptide and a gene for blasticidin deaminase were added downstream of the stop codon to enable selection for integration into the native *Pfstart1* locus. The gene construct was capped at the 3’ end with a second recombination block formed from the 3’UTR of *Pfstart1*. After CRISPR/Cas9-mediated transfection, integration into the *Pfstart1* locus in wild-type 3D7 parasites was selected for with blasticidin S and the transfected parasites were analysed by PCR which confirmed integration into the *Pfstart1* locus (Fig 1B). As a comparator, the PV-localised protein *Pf*PV1 (PF3D7_1129100)^20^ was C-terminally tagged with NG using CRISPR/Cas9 (S1A Fig). Correct integration into the *Pfpv1* locus was also verified by PCR (S1B Fig) and live cell microscopy confirmed the expected PV localisation (S1C Fig).

**Figure 1.**
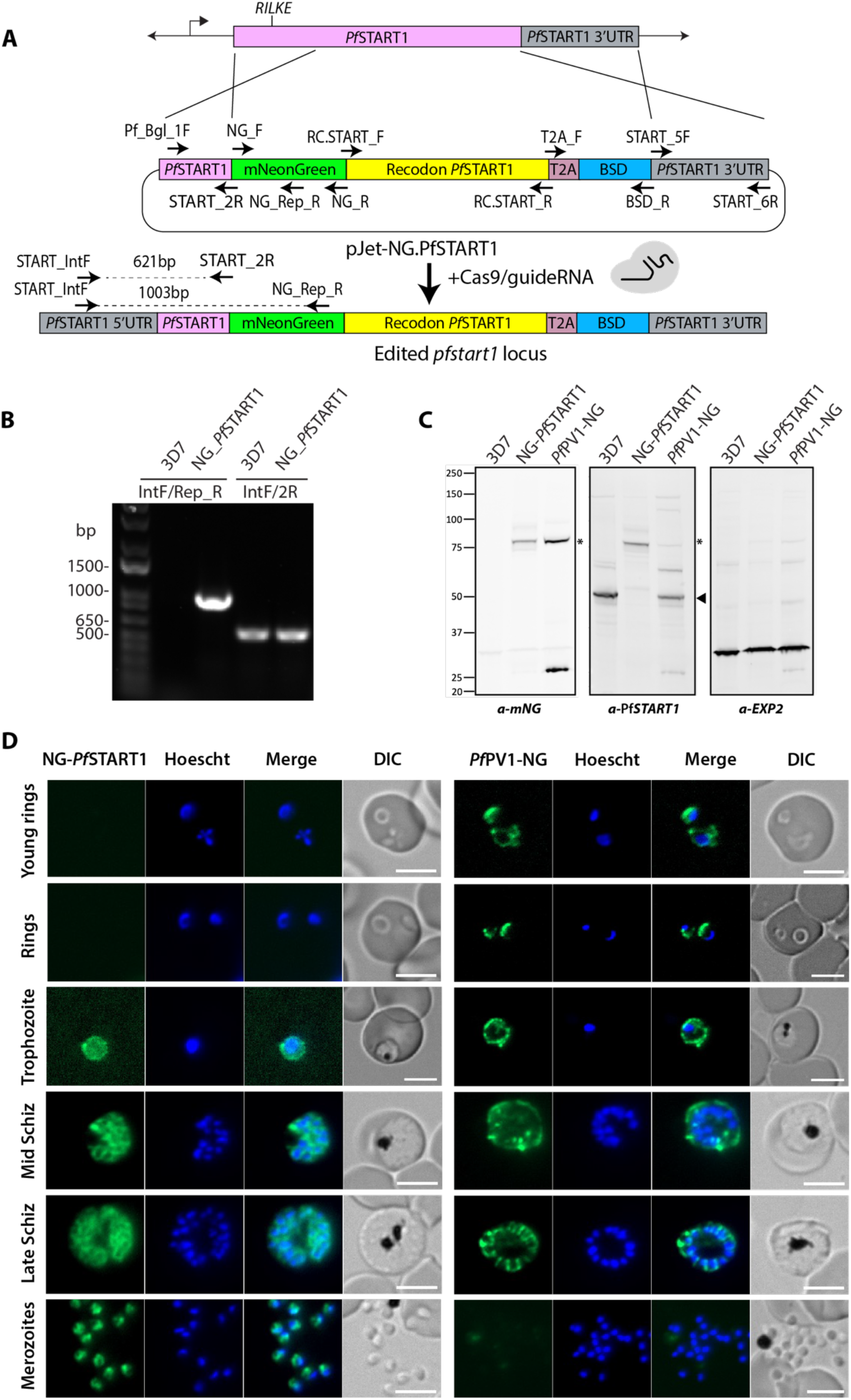
The mNeonGreen-*Pf*START1 fusion protein is strongly expressed in merozoites. **A)** *Pf*START1 was N-terminally tagged with mNeonGreen (NG) using recombination blocks derived from the 5’ coding region and 3’ UTR of *Pfstart1*. The START domain-containing region of *Pfstart1* was replaced by a recodonised sequence and integration was selected by exposing the parasites to blasticidin S. **B)** PCRs were performed on NG-*Pf*START1 parasites and parental 3D7 parasites with primers PfSTART_IntF (IntF), which primes upstream of the donor plasmid, and NG_Rep_R (Rep_R), which primes within the donor plasmid. These produced a product of the expected size in NG-*Pf*START1 parasites, indicating the locus was edited, but not in 3D7. Control primers IntF and START_2R (2R) produced a product of the expected size in both parasites. **C)** Western blot of mixed stage parasites with anti-NG IgG indicated the NG-*Pf*START1 parasites expressed a fusion protein of the expected size (∼75 kDa, asterisk). Parasites in which PV1 had been tagged with NG served as a positive control and 3D7, which lack NG, as a negative control. A ∼27 kDa anti-NG band is present in the *Pf*PV1-NG parasites, likely resulting from NG degradation from the fusion protein. A *Pf*START1-specific rabbit antibody also labelled the NG-*Pf*START1 fusion protein (asterisk) as well as the unmodified *Pf*START1 in the other parasites lines (∼48 kDa, arrowhead). The blots were also probed with anti-EXP2, which served as a loading control. **D)** Live cell imaging of the NG-*Pf*START1 asexual blood stage parasites indicated the fusion protein could be detected within trophozoites and schizonts and formed bright foci in merozoites. In contrast, *Pf*PV1-NG localised to the PV and was lost upon merozoite egress. Scale bars = 5 µm.

Western blot analysis of NG-*Pf*START1 and *Pf*PV1-NG parasite samples probed with anti-NG IgG labelled the ∼75 kDa NG-*Pf*START1 protein (PEXEL-cleaved *Pf*START1 including the T2A skip peptide is 48 kDa and NG is 27 kDa) or the *Pf*PV1-NG protein (predicted size ∼78 kDa) (Fig 1C). The NG-*Pf*START1 fusion protein was also labelled with a *Pf*START1-specific antibody ^10^ which labelled the smaller, non-tagged native *Pf*START1 in 3D7 parasites and *Pf*PV1-NG parasites (Fig 1C).

Microscopic analysis of live ring-stage parasites revealed *Pf*PV1-NG in the PV, whereas NG-*Pf*START1 was not detectable at this stage (Fig 1D). In trophozoites, the NG-*Pf*START1 protein was detected at the parasite periphery, like *Pf*PV1-NG, and was also expressed throughout the parasite and PV (Fig 1D). In mid and late schizonts, NG-*Pf*START1 was expressed throughout the cytoplasm of the merozoites and appeared to concentrate at the periphery. Following egress, the fusion protein remained detectable throughout the merozoite cytoplasm. In the majority of merozoites, the signal was not evenly spread but rather became concentrated in a region of the merozoite that did not colocalise with the nucleus (Fig 1D). In contrast to NG-*Pf*START1, *Pf*PV1-NG appeared to localise to the PV around the developing merozoites in late schizonts before being lost upon merozoite egress (Fig 1D).

As the lack of NG-*Pf*START1 signal in ring stages was unexpected, we performed immunofluorescence microscopy on previously generated *Pf*START1-HA parasites^10^. Again, we detected no signal in ring-stage parasites but observed cytoplasmic *Pf*START1-HA in trophozoites and in schizonts (S2 Fig) as seen with the NG-*Pf*START1 parasites.

### *Pf*START1 is released into the PV after merozoite invasion

As *PfS*TART1 has a PEXEL motif and other PEXEL proteins, such as the ring-infected erythrocyte surface antigen (RESA), have been shown to localise to the dense granules in merozoites^21^, we sought to determine if the region of concentrated NG-*Pf*START1 within the merozoite colocalised with the dense granule marker EXP2. As merozoites are small cells only 1-1.5 µm in length, expansion microscopy was attempted to better resolve the localisation of *Pf*START1 relative to markers of other secretory compartments^22^. Our attempts to detect NG-*Pf*START1 by expansion microscopy using commercial anti-NG IgGs were, however, not successful due to a weak signal which was difficult to discriminate from background labelling. We therefore probed *PfS*TART1-HA parasites we had previously prepared^10^ and in late schizonts observed a collection of several closely spaced small puncta within the merozoites (Fig 2A), which had not been distinguishable by lower-resolution live cell microscopy (Fig 1D). These puncta did not resemble or overlap with the rhoptry bulb marker RAP1, the dense granule marker EXP2 or the microneme marker AMA1^23–25^ (Fig 2A). This lack of colocalisation with merozoite organelles is consistent with previous observations of untagged *Pf*START1 in schizonts^8^. Having detected NG-*Pf*START1 by live cell microscopy in extracellular merozoites and in trophozoites and schizonts, but not in ring-stage parasites (Fig 1D), we sought to analyse the trafficking of *Pf*START1 immediately after invasion in more detail using lattice light-sheet microscopy. We examined 35 newly invaded merozoites across two imaging sessions to determine if NG-*Pf*START1 was released into the PV during or after invasion. NG-*Pf*START1 merozoites stained with SPY650-DNA were observed invading RBCs labelled with the membrane stain PKH26. The time taken for the strongly fluorescent centralised NG-*Pf*START1 punctum to transform into several peripheral smaller foci around the merozoite, which colocalise with the nascent PV, was recorded (Fig 2B, S1 Video). The formation of these peripheral foci likely corresponds to the secretion of NG-*Pf*START1 into the nascent PV, which commenced, on average, 109 (s. d. 71) seconds after the merozoite had finished invading the RBC (Fig 2C). To examine when *Pf*START1 has been completely secreted into the nascent PV, the mean intensity of NG across 35 invasion events was measured after the merozoite had invaded the RBC by spot detection and normalised between 0-1. This analysis found that the mean NG-*Pf*START1 intensity begins to diminish soon after invasion (Fig 2D), which we interpret as the beginning of secretion of NG-*Pf*START1 into the nascent PV, in agreement with our above measurement (Fig 2C). Then, the mean NG-*Pf*START1 intensity plateaus at ∼850 seconds, which we interpret as the complete dispersal of NG-*Pf*START1 throughout the expanding PV (Fig 2D, S3 Table).

**Figure 2.**
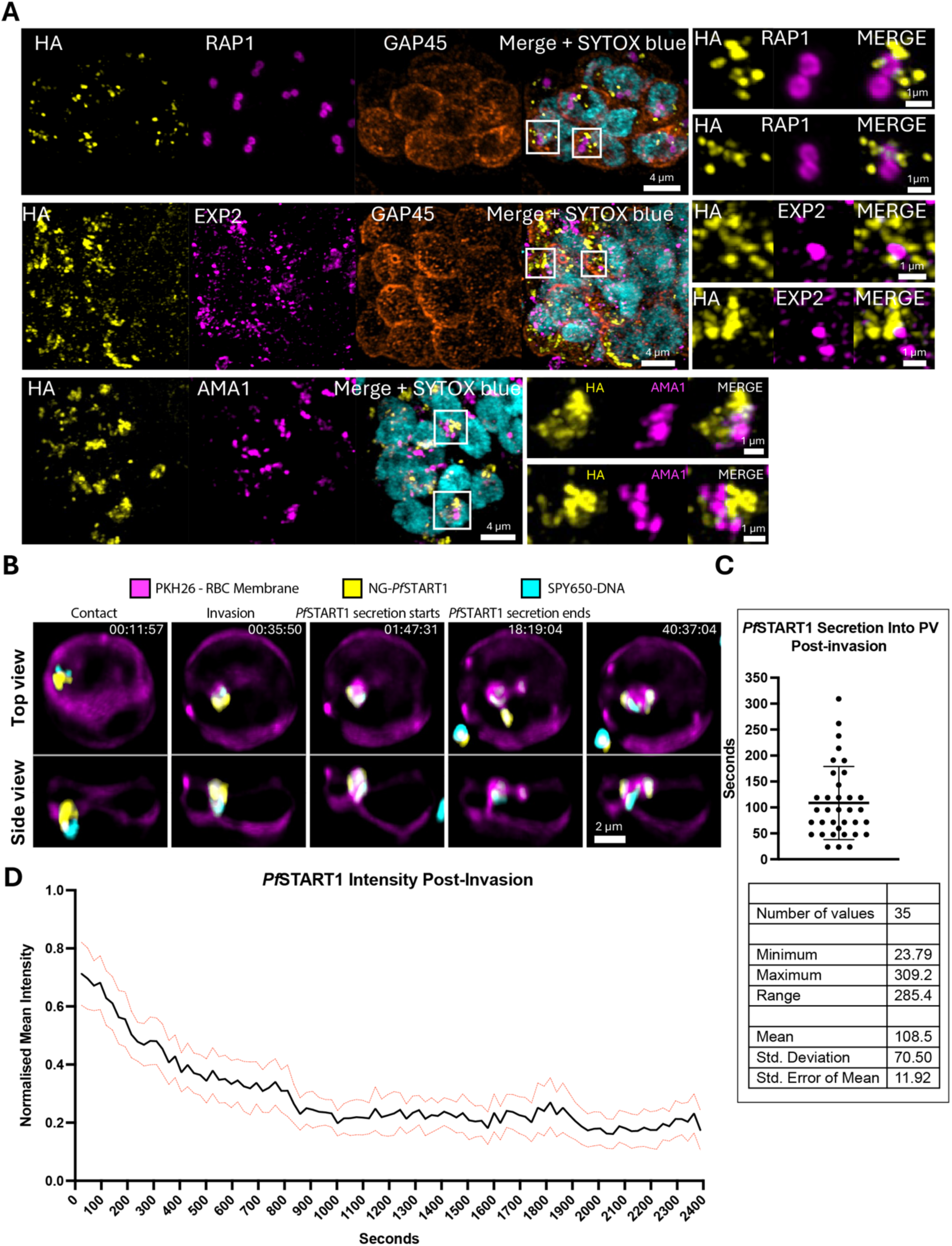
*Pf*START1 forms numerous foci with the merozoite and appears to be secreted into the PV on average 10G seconds after the merozoite invades the RBC. **A**) Confocal maximum intensity projections of expanded *Pf*START1-HA (B4 clone) schizonts illustrating that *Pf*START1-HA localises to several clustered foci within the merozoite that do not colocalise with markers of the rhoptry bulb (RAP1), dense granules (EXP2) or micronemes (AMA1). GAP45 labels the merozoite periphery and SYTOX™ blue stains DNA. The white boxes in the merged image indicate zoomed areas to the right. **B)** Live-cell imaging of NG-*Pf*START1 schizonts stained with SPY650-DNA (DNA marker) and RBCs stained with PKH26 (RBC membrane marker) was performed using lattice light-sheet microscopy. 35 invasion events across two imaging sessions were observed, with a representative image series shown. Frames demonstrate that NG-*Pf*START1 is secreted into the PV ∼107 seconds after invasion, subsequently colocalising with the expanding PV. **C)** On average, NG-*Pf*START1 is secreted into the PV ∼108.5 s after invasion is complete. Error bars represent the standard deviation of the mean of 35 invasion events measured across two imaging sessions. **D)** The mean intensity of NG-*Pf*START1 was measured after invasion of RBCs by spot detection in Imaris, and normalised (min-max normalisation). Red dotted lines represent the 95% confidence interval of the mean (black line) of 35 invasion events measured in two imaging sessions.

### Structural analysis of *Pf*START1 indicates that WEHI-GG1 competes for the predicted lipid binding site and that the inhibitor cannot efficiently bind to *Pb*START1 due to key amino acid differences

Having shown that *Pf*START1 enters the PV after invasion, where its inhibition likely arrests PV expansion, we next investigated why the *Pf*START1 inhibitor WJM-715 had not prevented the growth of *P. berghei* parasites, which must invade and expand within mouse RBCs ^14^. To determine whether structural differences between *Pb*START1 and its *Pf*START1 orthologue might explain the failure of WJM-715 to reduce the parasitaemia in *P. berghei*-infected mice, we aligned these protein sequences as well as that of *P. knowlesi* - a species closely related to the other major human pathogen *P. vivax* (S3 Fig). Within the START domain, there are 30 amino acids predicted to line the phospholipid binding pocket, including three amino acids which, when mutated (I224F, N309K and N330K), are known to confer resistance to the *Pf*START1 inhibitor WEHI-991 ^10,14^. 21/30 of these amino acids are identical between the species (S3 Fig). Interestingly, compared to *P. falciparum*, four positions in *P. knowlesi* and nine positions in *P. berghei* are not identical, including position 330 (N in *P. falciparum* and *P. knowlesi* and S in *P. berghei*) which we have previously found to be involved in *Pf*START1-inhibitor resistance ^10^.

To understand how these amino acid differences might influence the potency of the compound against the different *Plasmodium* species we resolved the crystal structure of *Pf*START1 in the presence of WEHI-991 to a resolution of 1.6 Å (PDB ID: 44ZY / PDB_000044ZY) (Table S4). From this structure it can be seen that WEHI-991 occupies the central hydrophobic cavity of *Pf*START1 (Fig 3A). Notably, all three resistance-conferring residues - I224, N309 and N330 - contact WEHI-991. Specifically, I224 contacts the gem dimethyl group, whereas N309 and N330, together with Y328, interact with both the carboxamide N-H and the aryl amine N-H through a water-mediated hydrogen-bonding network (Fig S4). Additionally, N279 and N228 form hydrogen bonds with the sulfonamide group, while N228 also hydrogen bonds to the carboxamide carbonyl. Overall, the WEHI-991 binding cavity is largely neutral in charge – devoid of both basic and acidic amino acids (Fig S5). Substitution of I224 with Phe, or N309 and N330 with Lys, is predicted to disrupt ligand binding through steric clashes and altered hydrogen-bonding interactions, providing a molecular explanation for resistance.

**Figure 3.**
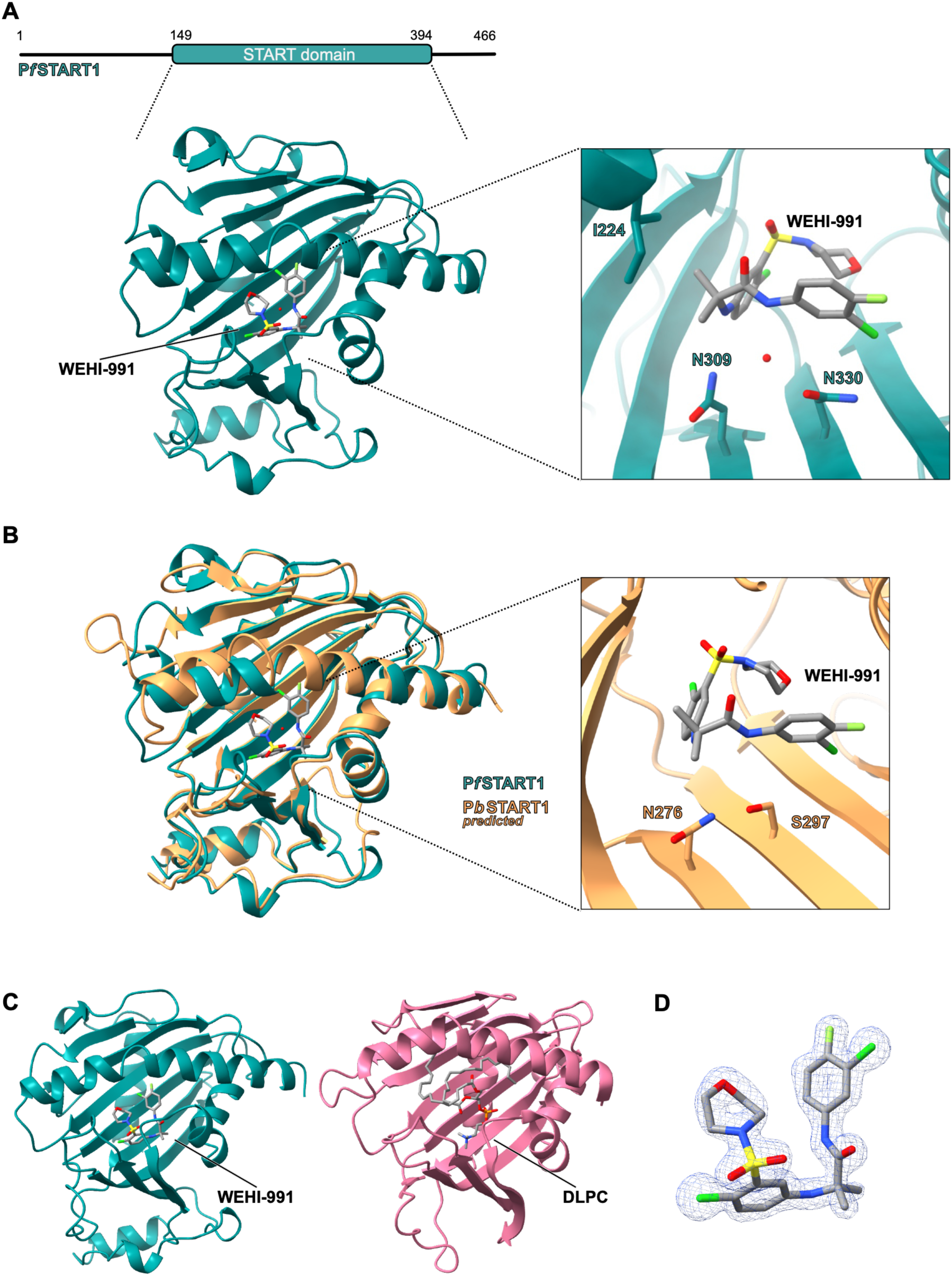
Structural analysis demonstrates that WEHI-GG1 occupies the central lipid binding cavity of *Pf*START1. **A)** (top) The domain architecture of *Pf*START1 (teal). A construct of residues 149-394 were amenable to crystallisation. (bottom) The crystal structure of *Pf*START1 (teal) in complex with WEHI-991 solved to 1.6 Å. Callout: the ligand-binding pocket; key residues N309 and N330 are highlighted. **B)** Overlay of the crystal structure of *Pf*START1 (teal) with WEHI-911 with the AlphaFold3-predicted structure of *Pb*START1 (orange). Callout: the ligand-binding pocket of the predicted structure of *Pb*START1 overlayed with WEHI-991. The equivalent residues to *Pf*START1 N309 (N276) and N330 (S297) are highlighted. **C)** The crystal structure of *Pf*START1 (teal) with WEHI-991 juxtaposed with the crystal structure of human phosphatidylcholine transfer protein (PC-TP) START domain (pink) with 1,2-dilinoleoyl-*sn*-glycerol-3-phosphorylcholine (DLPC, PDB: 1LN2). **D)** A 1.6 Å composite omit difference map for WEHI-991, together with the associated omit density displayed as a semi-transparent blue surface contoured at 2.0 σ.

To structurally compare *Pf*START1 and *Pb*START1 we modelled *Pb*START1 using AlphaFold3. The START domain of *Pb*START1 aligned well with our crystal structure of *Pf*START1 (Fig 3B, r.m.s.d. 0.8 Å). Although most residues within the binding pocket were conserved, the desensitising residue S297 in *Pb*START1 occupied the same position as N330 in *Pf*START1. Given that N330 participates in a water-mediated hydrogen-bonding network with WEHI-991, substitution of this residue with Ser is predicted to weaken ligand binding, providing a molecular explanation for the reduced activity of these compounds against *P. berghei* and in mouse models.

Phosphatidylcholine has previously been shown to bind to recombinant *Pf*START1 ^13^. To investigate this further we compared our crystal structure with the crystal structure of the human phosphatidylcholine transfer protein (PC-TP) START domain (Fig 3C, PDB: 1LN2) ^26^. PC-TP was crystalised with the natural polyunsaturated phospholipid 1,2-dilinoleoyl-*sn*-glycerol-3-phosphorylcholine (DLPC) present in the central cavity. By overlaying the structures, we observed that WEHI-991 and DLPC both occupy the same cavity, strongly suggesting the compound is a competitive inhibitor of phospholipid binding (Fig S4B). Interestingly, it appears that DLPC occupies considerably more space than WEHI-991. We determined the volume of the accessible central cavity of *Pf*START1 to be 748.44 Å^3^; given that WEHI-991 only occupies 346 Å^3^ (Fig 3D), this suggests there is still more physical space for compound optimisation (S4C Fig). Indeed, this is corroborated by the relatively few points of interaction between the WEHI-991 difluorobenzene ring and the surface of the cavity (Fig S4A).

### *P. falciparum* parasites in which the START domain has been replaced with an orthologous region from *P. berghei* are viable

To confirm that the amino acid differences between *Pf*START1 and *Pb*START1 prevent the inhibitors from efficiently targeting *Pb*START1, we replaced the START domain of *Pf*START1 with the equivalent domain from *Pb*START1 by homologous recombination (Fig 4). To target the *Pfstart1* locus, we constructed a donor plasmid containing 5’ and 3’ homology blocks from *Pfstart1* bordering the START domain of *Pb*START1 (Fig 4A) and a hDHFR resistance marker.

**Figure 4.**
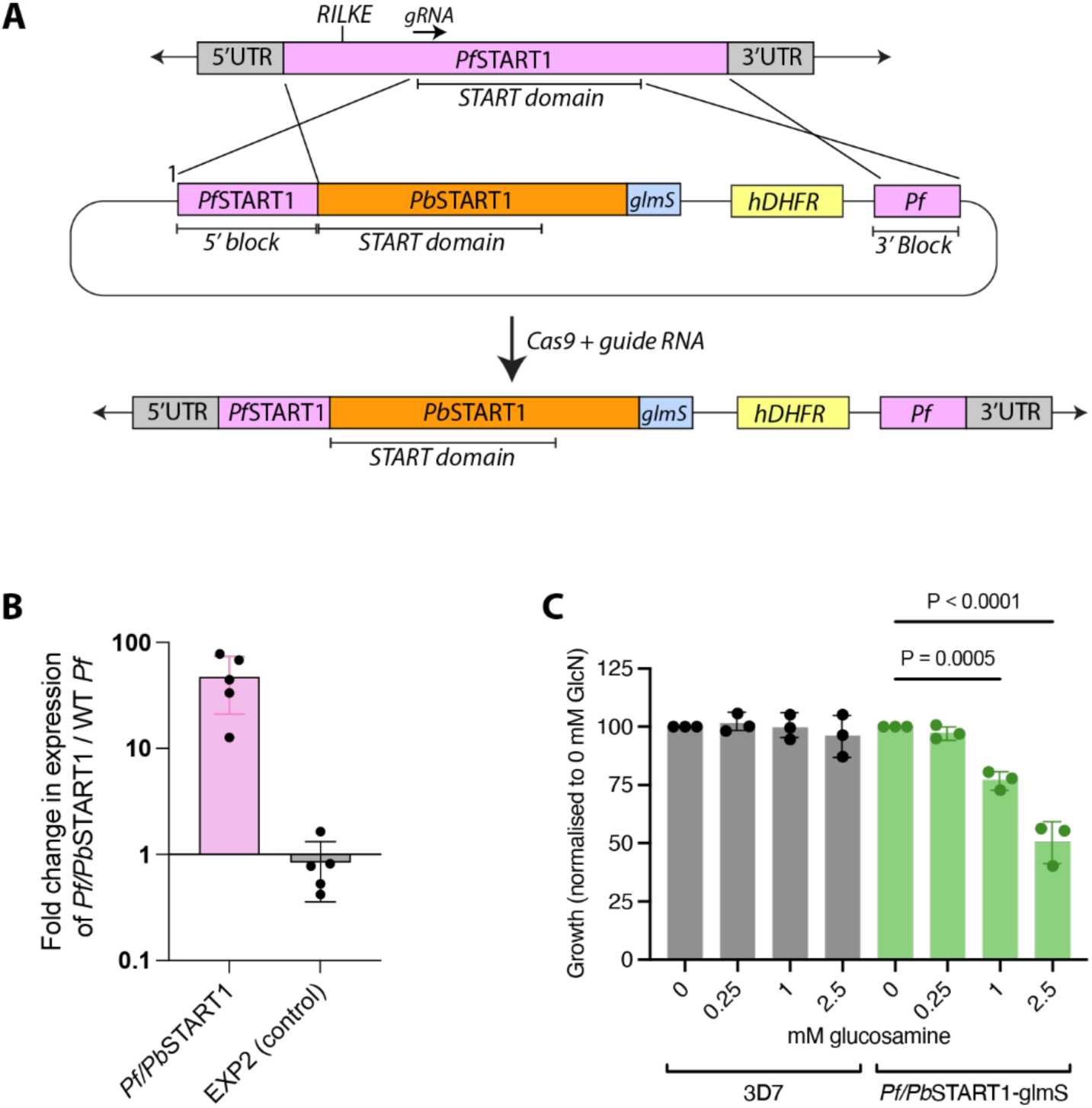
*P. falciparum* parasites in which the START domain of *Pf*START1 has been replaced with an orthologous region from *P. berghei* are viable. **A**) The START domain and C-terminal region of *Pf*START1 were replaced with orthologous regions from *Pb*START1 using *P. falciparum* 5’ and 3’ homology blocks. Cas9 complexed with a *Pf*START1-specific guide RNA was co-transfected with the donor plasmid to facilitate the gene replacement event. Primer pairs used to validate correct integration into the *Pfstart1* locus are shown in S2 Table and on S4 Fig gene maps. **B)** RT-qPCR with primers specific for the *Pf/pbstart1* gene amplified a product that was ∼50 times more abundant in these parasites than in the 3D7 parental line. Control RT-qPCR with *exp2* primers amplified a similarly abundant product in both parasite lines. **C)** To determine if *Pf*/*Pb*START1 is crucial for blood-stage growth, its expression was knocked down in non-clonal parasites by activating the *glmS* riboswitch with glucosamine (GlcN) to degrade the gene’s mRNA. Parasites were treated with GlcN for 72 hours (1.5 intraerythrocytic growth cycles) and their growth was quantified by measuring levels of parasite lactate dehydrogenase (LDH) activity. LDH activity was normalised to the 0 mM GlcN condition, representing 100% growth. Wild-type 3D7 parasites containing no *glmS* riboswitch were employed as a control to demonstrate that GlcN was not significantly reducing growth. Points and error bars indicate the mean and standard deviation of three independent experiments, each conducted in technical triplicate. For each parasite line, the 0 mM GlcN condition was compared to the 0.25, 1 and 2.5 mM conditions using one-way ANOVA with Šidák’s multiple comparisons test. No bar = not significant.

After selecting for transgenic parasites using WR99210, we performed diagnostic PCRs with primer pairs outside and inside the donor plasmid. These PCRs indicated that the *P. falciparum* START domain had been replaced with the *P. berghei* domain (S6A, B Fig). As the chimeric protein lacked an epitope tag, to determine if it was expressed, RT-qPCR was performed with a primer set specific for the chimeric mRNA. These primers amplified a product that was about 50 times more abundant in the *Pf/Pb*START1 parasites than in the parental WT *P. falciparum* 3D7 parasites (Fig 4B). In contrast, primers to the control *exp2* gene amplified a similarly abundant product in both parasite lines.

To establish that *Pf/Pb*START1 was functional and important for parasite growth, like the endogenous protein, the expression of the chimeric protein was knocked down by adding glucosamine (GlcN) to the *Pf/Pb*START1 parasites. After 72 hours’ continuous exposure to GlcN, growth was quantified by measuring parasite lactate dehydrogenase (LDH) activity. This showed that growth was unaffected at 0.25 mM GlcN but reduced at 1 mM and at 2.5 mM GlcN to ∼75% and ∼50% of growth in the absence of GlcN, respectively (Fig 4C). This is comparable to the ∼40% reduction in growth previously observed when *Pf*START1 was knocked down using 2.5 mM GlcN for 72 hours ^10^.

### *P. falciparum* parasites expressing the *Pf/pb*start1 gene are resistant to the *Pf*START1 inhibitors MMV006833, WEHI-GG1 and WJM-715

Next, we challenged *Pf/Pb*START1 parasites with the aryl acetamide *Pf*START1 inhibitor MMV006833 (M-833) and its potent analogues WEHI-991 and WJM-715 for 72 hours (S1 Table) ^10,14^. LDH activity was then measured and the EC_50_s of the compounds were found to be ∼150-600-fold higher in *Pf/Pb*START1 parasites than in parental 3D7 parasites (Fig 5).

**Figure 5.**
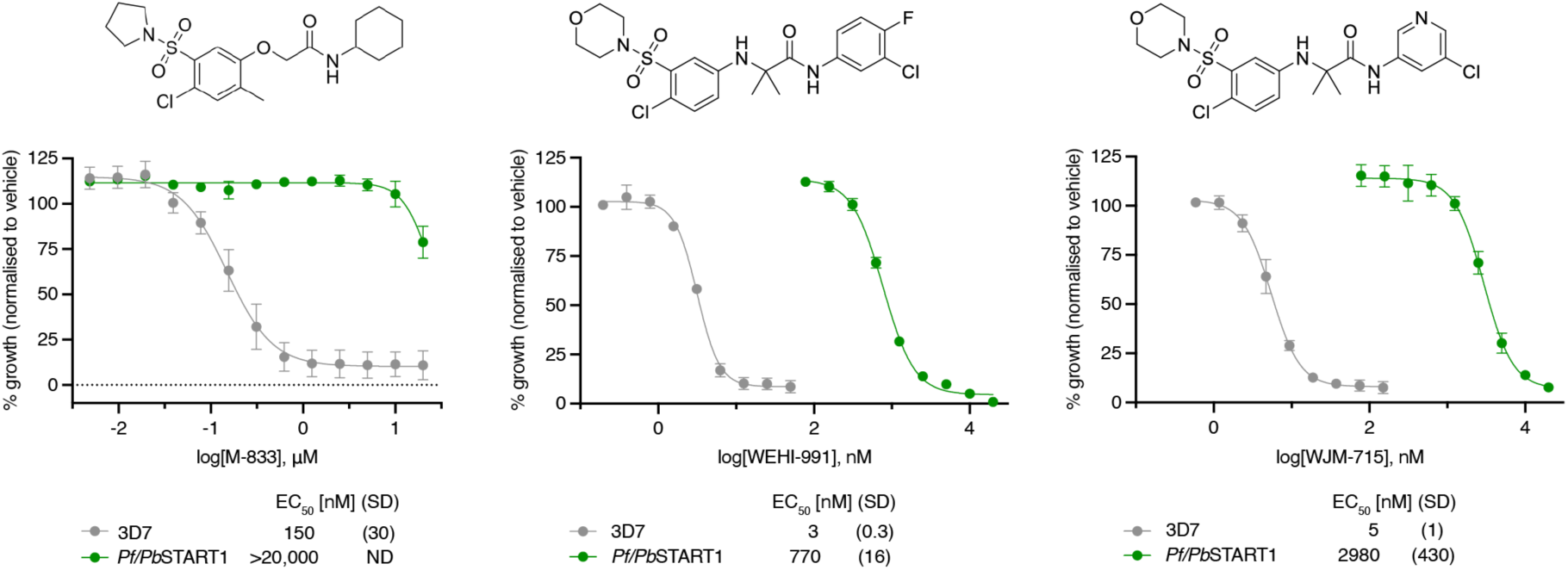
*P. falciparum* parasites expressing the *Pf/pb*start1 gene are highly resistant to the *Pf*START1 inhibitors M-833, WEHI-GG1 and WJM-715. **A**) *Pf/Pb*START1 (green) and parental 3D7 (grey) parasites were treated with serially diluted inhibitors for 72 hours. Growth was quantified by measuring lactate dehydrogenase activity in lysed parasite samples and normalised to growth in the presence of the DMSO vehicle control. Points and error bars represent the mean and standard deviation of three independent experiments, each conducted in technical duplicate or triplicate. Mean EC50 values (µM) and standard deviations are indicated; n=3.

The above data led us to speculate that the failure of WJM-715 to suppress the parasitaemia in *P. berghei*-infected mice was almost certainly due to the *Pf*START1 inhibitor’s inability to bind *Pb*START1. We then wondered if any other *Pf*START1-inhibitory analogues of WEHI-991 would inhibit *Pb*START1 and might therefore be more effective against murine malaria than WJM-715. To investigate this, we tested 11 previously-synthesised analogues of WEHI-991 against *Pf/Pb*START1 parasites and 3D7 parental parasites ^14^. The analogues’ activities were measured in 72-hour growth assays using a three- or four-point dilution series of each compound, with a maximum concentration high enough to completely suppress 3D7 parasite growth. Eight of the compounds demonstrated virtually no activity against *Pf/Pb*START1 parasites compared to strong inhibition of the parental 3D7 parasites (Fig 6A, triangle versus circle symbols, respectively). Three compounds did show some activity against *Pf/Pb*START1 parasites but were still more inhibitory against 3D7 parasites (Fig 6B, triangle versus circle symbols, respectively). To more accurately measure the potency of the three inhibitory compounds against *Pf/Pb*START1, growth assays were performed with a full nine-point dilution series of the inhibitors. Where EC_50_ values could be determined, the analogues were ∼5-10-fold more potent against 3D7 than *Pf/Pb*START1 parasites. Interestingly, 3D7 parasites were substantially less sensitive to **10y** (EC_50_ 0.416 µM) than to WJM-715 (EC_50_ 0.005 µM), whereas *Pf/Pb*START1 parasites were more sensitive to **10y** (EC_50_ 2.13 µM) than to WJM-715 (EC_50_ 2.98 µM). Despite this, and although the fold change in EC_50_ between the lines was less than for M-833, WEHI-991 and WJM-715, the analogues **10y**, **13i** and **13h** were overall no more active against *Pf/Pb*START1 parasites than were WEHI-991 and WJM-715 (Fig 6C) and would therefore be similarly unsuitable for use in mouse studies.

**Figure 6.**
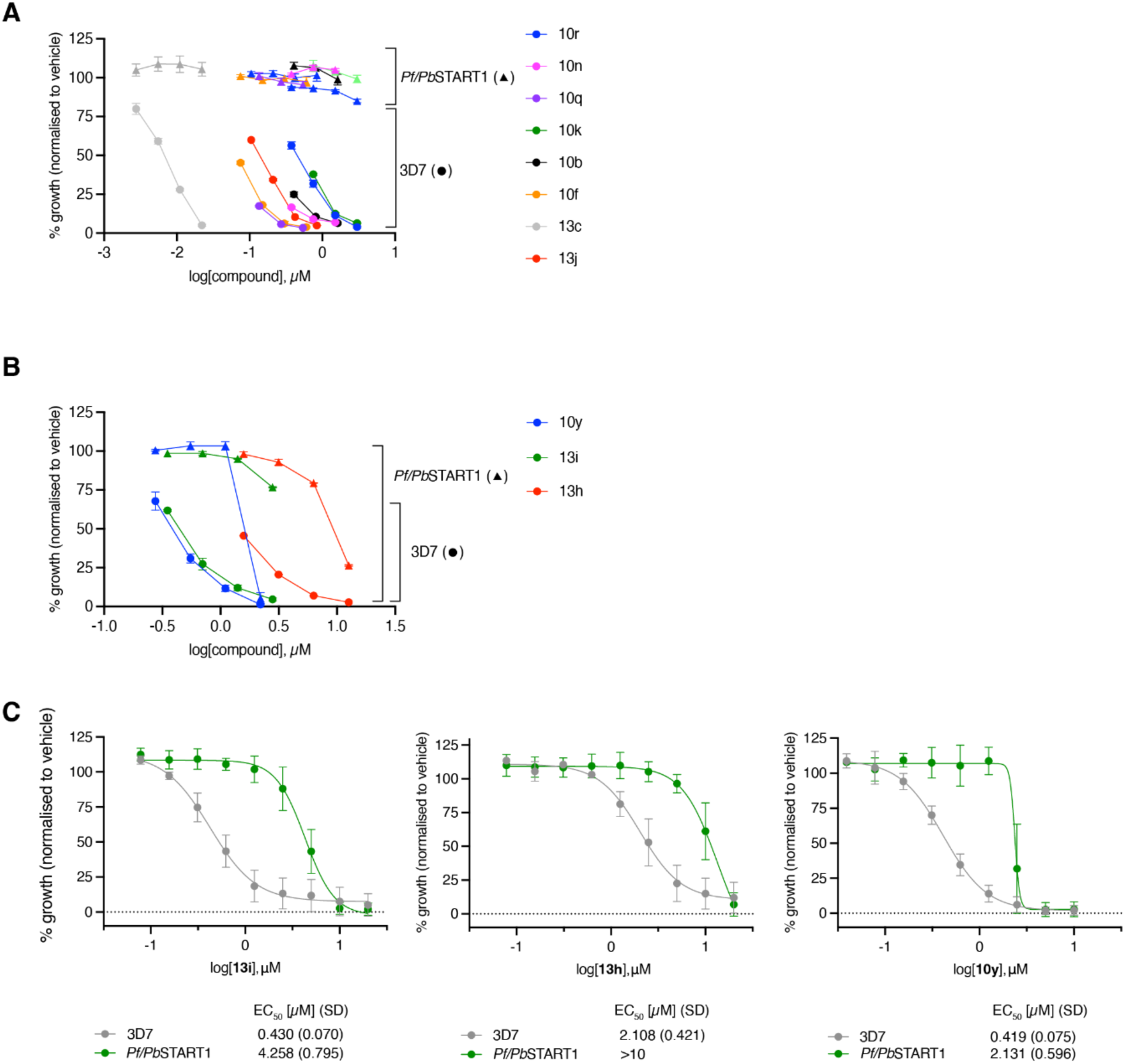
Transfectant *P. falciparum* parasites expressing the *Pf/pbstart1* gene show partial sensitivity to some *Pf*START1 inhibitors. **A)** In a 72-hour growth assay, a series of *Pf*START1-targeted analogues were tested against chimeric (*Pf/Pb*START1; triangle symbols) and wild-type (3D7; circle symbols) parasites. The maximum concentration of each compound corresponded to approximately double the EC50 value as determined in 72-hour growth assays using 3D7 parasites ^14^. At the concentrations used, the analogues demonstrated little to no activity **(A)** or partial activity **(B)** against *Pf/Pb*START1 parasites. Points and error bars represent the mean and standard deviation of technical duplicates. **C)** To assess the potency of the compounds with partial activity against the chimeric parasite line, parasites were treated with a dilution series of each compound for 72 hours. The potency was greatly reduced in the chimeric parasite line. Points and error bars represent the mean and standard deviation of three independent experiments, each conducted in technical duplicate. Mean EC50 values (µM) and standard deviations are indicated; n=3.

### *Pf*START1 inhibitors block *Pf/Pb*START1 parasite invasion of RBCs but not as potently as they block parental 3D7 parasite invasion

As *Pf*START1 inhibitors act primarily by blocking wild-type parasite invasion of RBCs, we sought to confirm that the inhibitors have the same mechanism of action against *Pf/Pb*START1 parasites. Egress and invasion assays were first performed on 3D7 parasites expressing an exported nanoluciferase (Hyp1-Nluc) reporter ^9,27^. Late-stage schizonts were incubated with the inhibitors for 4 hours during which time they underwent egress, releasing the Hyp1-Nluc reporter into the culture medium. Across a range of concentrations, the inhibitors did not reduce the extracellular bioluminescence signal relative to the DMSO vehicle control, indicating that they do not inhibit egress (Fig 7A). After the 4-hour treatment window, compounds and unruptured schizonts were removed. The newly invaded parasites were incubated until they reached the trophozoite stage (24 hours), when Hyp1-Nluc is expressed, and bioluminescence levels were then measured in parasite lysates. A concentration-dependent reduction in bioluminescence occurred, indicating that the *Pf*START1 inhibitors blocked invasion as determined previously for M-833 (Fig 7A) ^9^. As the *Pf/Pb*START1 parasites do not express Hyp1-Nluc, the levels of LDH in parasite lysates were measured 24 hours after the 4-hour treatment window. This revealed that the inhibitors blocked *Pf/Pb*START1 parasite invasion of RBCs, but only at high concentrations (Fig 7B).

**Figure 7.**
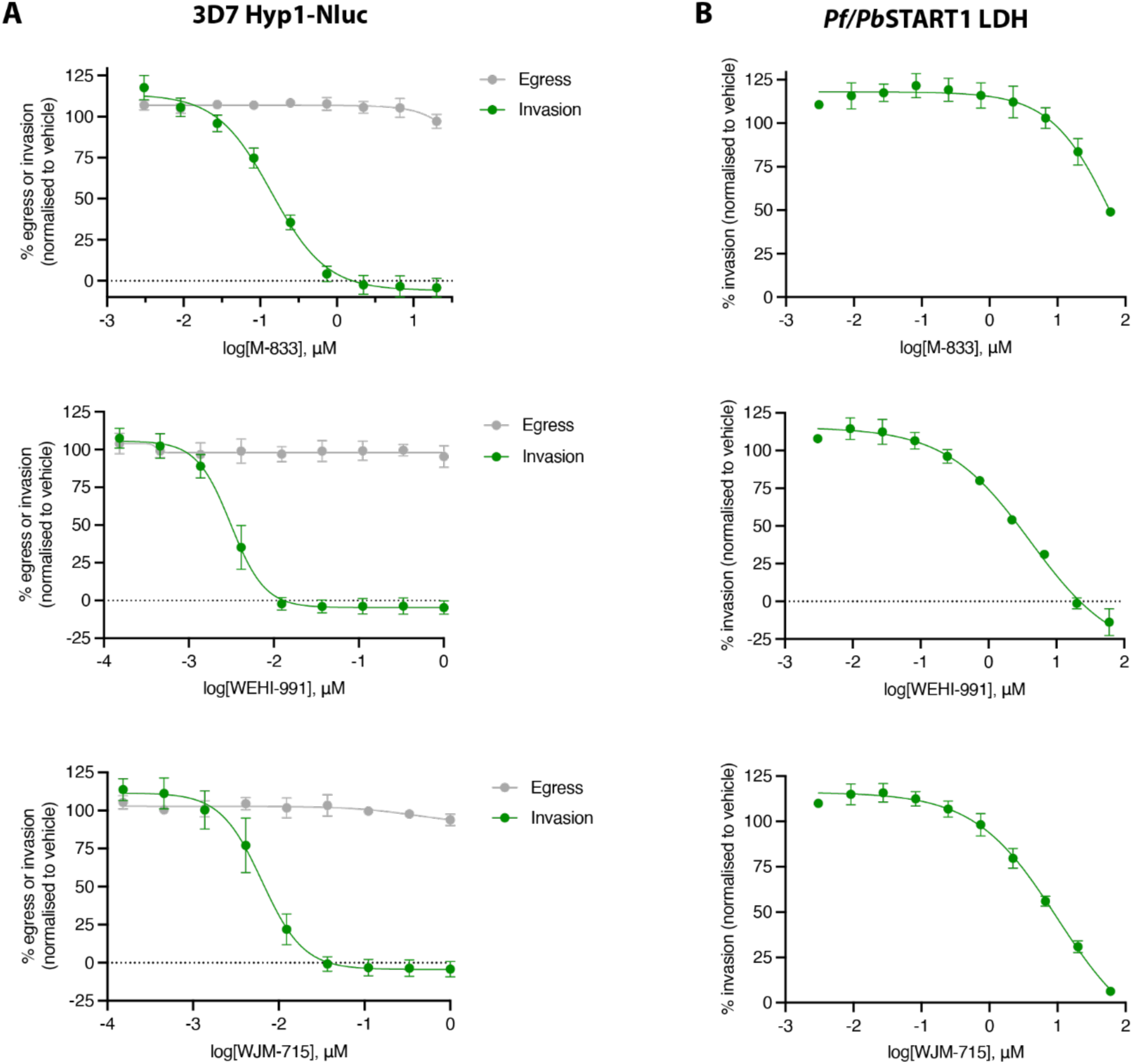
*Pf*START1 inhibitors block *Pf/Pb*START1 parasite invasion of RBCs but less potently than parental 3D7 parasite invasion. **A)** Egress and invasion assays were performed on 3D7 parasites expressing the exported bioluminescent protein Hyp1-Nluc, treated with *Pf*START1 inhibitors M-833, WEHI-991 and WJM-715. In this assay, late-stage schizonts were incubated with the compounds for 4 hours, during which time egress and invasion occurred. To quantify egress, the bioluminescence of Nluc released into the medium during the incubation period was measured. This indicated that none of the compounds inhibited egress. After 24 hours, the bioluminescence of trophozoite-stage parasites that had invaded during the drug-treatment window was measured and indicated that the compounds strongly inhibited invasion. **B)** *Pf/Pb*START1 parasites were similarly treated but as they do not express Hyp1-Nluc the degree of invasion could only be ascertained by measuring trophozoite LDH levels. This indicated that the inhibitory compounds also reduced *Pf/Pb*START1 parasite invasion, but higher concentrations were required to achieve this compared to the 3D7 Hyp1-Nluc parasites.

### *P. knowlesi* remains susceptible to the *Pf*START1 inhibitors

Having found that the *Pf/Pb*START1 parasites were resistant to the *Pf*START1 inhibitor series, we next tested the compounds against *P. knowlesi* parasites, which share more conserved residues in the predicted lipid binding pocket with *P. falciparum* than does *P. berghei* (S3 Fig). *P. knowlesi* parasites adapted to grow in human normocytes ^28^ were cultured in the presence of inhibitors for 1.5 cell cycles (48 hours) and their growth was measured using a Sybr green DNA stain assay (S7 Fig). Overall, the *P. knowlesi* parasites were slightly less susceptible to the compounds than *P. falciparum* 3D7 parasites but were not nearly as resistant as the *Pf/Pb*START1 parasites (Table 1). The exception to this was compound WJM-715, to which *P. knowlesi* was about five times more sensitive than *P. falciparum*.

**Table 1.** Sensitivity of *P. knowlesi* parasites to M-833 compound series compared to *P. falciparum* 3D7 and *Pf/Pb*START1 parasites. NA – Not available.

| Compound | *Pf* 3D7 (72 h) |  | *Pf/Pb*START1 (72 h) |  | *Pk* YH1 WT (48 h) |  |
| --- | --- | --- | --- | --- | --- | --- |
|  | EC<sub>50</sub> (μM) | SD | EC<sub>50</sub> (μM) | SD | EC<sub>50</sub> (μM) | SD |
| **M-833** | 0.155 | 0.032 | >20 | NA | 2.20 | 1.50 |
| **WEHI-991** | 0.003 | 0.0003 | 0.770 | 0.016 | 0.05 | 0.053 |
| **WJM-715** | 0.005 | 0.001 | 2.984 | 0.430 | 0.001 | 0.001 |
| **13i** | 0.430 | 0.070 | 4.258 | 0.795 | NA | NA |
| **13h** | 2.108 | 0.421 | >10 | NA | NA | NA |
| **10y** | 0.419 | 0.075 | 2.131 | 0.596 | NA | NA |

### WJM-715 does not clear the parasitaemia of mice infected with chimeric *P. berghei* parasites expressing a *Pf*START1 protein

Since we could not find WEHI-991 analogues that were highly potent against *P. falciparum* parasites expressing a *P. berghei* START domain, we generated chimeric *P. berghei* parasites expressing a *P. falciparum* START domain and challenged these *P. berghei Pb/Pf*START1 parasites *in vivo* with the *P. falciparum*-specific inhibitors. A CRISPR/Cas9 approach was used to replace the START domain of *Pb*START1 in the *P. berghei* parasites with the equivalent region from *P. falciparum* (Fig 8A and S8A Fig). PCRs performed with genomic DNA from the transfected *Pb/Pf*START1 parasites indicated the domain had been successfully replaced (S8B Fig). Western blots of these chimeric parasites with an antibody specific for *Pf*START1 indicated the presence of the *P. falciparum* domain, which was absent from the parental *P. berghei* parasites (Fig 8B). The full-length fusion protein was predicted to be 50 kDa and the plasmepsin V cleaved form, 44 kDa. The observed doublet bands labelled with the *Pf*START1-specific antibody were smaller than this suggesting there could be further processing as previously observed for *Pf*START1 (Fig 8B) ^10^.

**Figure 8.**
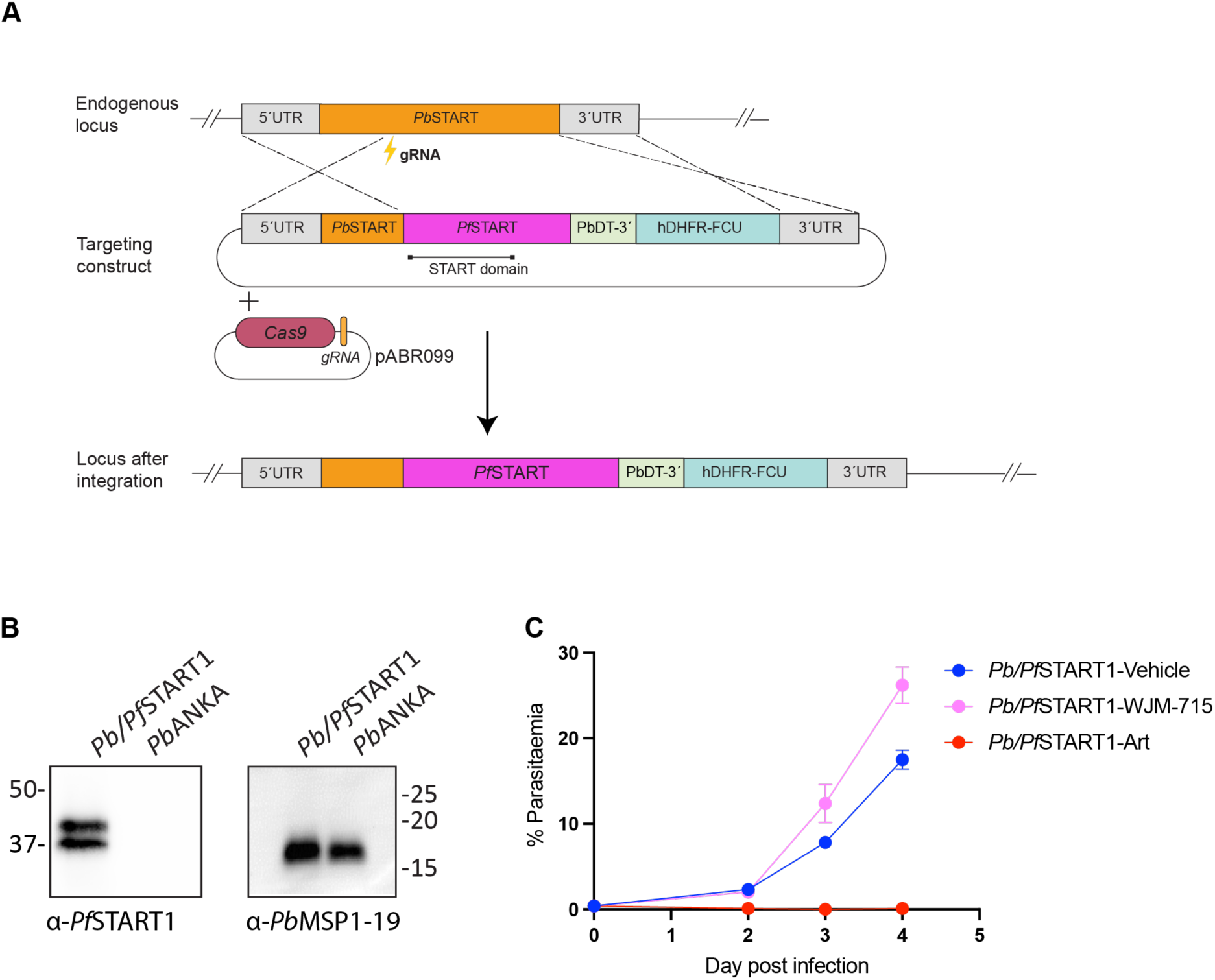
Replacement of the START domain in *P. berghei* parasites with that from *Pf*START1 did not facilitate parasite clearance with the *P. falciparum* inhibitor WJM-715. **A**) The START domain of *Pb*START1 was replaced with that of *Pf*START1 by homologous recombination following Cas9 cleavage of the *Pbstart1* gene. The donor plasmid was made using 5’ and 3’ homology blocks containing the 5’UTR/coding sequence and 3’UTR of *Pbstart1* to flank the *Pfstart1* replacement gene. A hDHFR-FCU gene cassette was also included to select for integration using pyrimethamine. Gene replacement was facilitated by co-transfection with a pABR099 plasmid, which encodes the Cas9 enzyme and a START1-specific guide RNA. **B)** Western blot analysis of *P. berghei* wild-type (*Pb*ANKA) and *Pb/Pf*START1 parasites confirmed correct replacement and expression of the *Pbstart1* gene as detected with a *PfS*TART1-specific rabbit antibody. A *Pb*MSP1-19-specific antibody served as a loading control and was detected in both parasite lines. **C)** Oral administration of WJM-715 at 50 mg/kg failed to suppress the parasitaemia in mice infected with *Pb/Pf*START1 parasites, resulting in an increase in parasitaemia like that of mice treated with the DMSO vehicle control. In contrast, artesunate (Art) successfully prevented an increase in parasitaemia.

To test the efficacy of WJM-715, the compound was administered orally at 50 mg/kg to 6-week-old Swiss mice that had recently been infected with the chimeric *Pb/Pf*START1 parasites (n=4). A DMSO vehicle control was also administered (n=4) as well as the known parasite growth suppressor artesunate at 30 mg/kg (n=1). The dosage regime was repeated at 1, 2 and 3 days post-infection with parasite growth being monitored by microscopic observation of blood drawn from the tail. We observed that WJM-715 did not supress parasite growth, resembling the vehicle control, whereas artesunate strongly suppressed parasite growth (Fig 8C).

## Discussion

*Pf*START1 possesses a typical PEXEL leader sequence with a recessed hydrophobic signal sequence and a PEXEL-like motif, RILKE, which is cleaved by the ER-resident protease plasmepsin V ^8^. This, however, does not result in export of the protein into the RBC compartment, possibly due to the P_1_’ lysine (4^th^ position in the PEXEL motif) inhibiting the protein’s interaction with the protein export translocon PTEX ^8^. In merozoites, PTEX components and exported proteins such as RESA have been observed in the dense granule organelles, from which they are released into the PV after invasion ^29,30^. To resolve the location of *Pf*START1 in live parasites we tagged the protein near its PEXEL-processed N-terminus with NG and observed parasites by widefield fluorescence microscopy. NG-*Pf*START1 was first visible in trophozoites where it appeared to be concentrated within internal structures and at the parasite periphery, possibly corresponding to the PV. In contrast, the secreted *Pf*PV1-NG protein strongly concentrated in the PV, with a typical “necklace of beads” appearance ^20^. NG-*Pf*START1 was strongly expressed in schizonts and was retained in free merozoites after egress, unlike the *Pf*PV1-NG signal which dispersed after breakdown of the PV. In most ring-stage parasites, as defined by their amoeboid shape, we did not observe any NG-*Pf*START1 fluorescence. To ensure the N-terminal tagging of *Pf*START1 with NG had not changed the timing of protein expression, we performed immunofluorescence microscopy on *Pf*START1-HA parasites and again observed no expression in ring-stage parasites.

To more closely examine the localisation of *Pf*START1 immediately after invasion, we imaged live, newly invaded NG-*Pf*START1 merozoites using the gentle lattice light-sheet method. We observed that NG-*Pf*START1 began to distribute from the interior of the merozoite to the merozoite circumference (presumably indicative of PV localisation) on average 109 (s. d. 71) seconds after invasion was complete. During this initial period, the fluorescence signal intensity within the merozoite declined, likely due to the secretion of NG-*Pf*START1 into the PV. The signal plateaued ∼850 seconds after merozoite internalisation, suggesting dispersal of the fluorescent protein in the PV was complete at this point. Interestingly, we did not observe NG-*Pf*START1 in amoeboid ring-stage parasites by live cell widefield microscopy, which suggests that the fusion protein present in merozoites does not persist for long in parasites post-invasion and must be re-synthesised during intracellular parasite development. In very young (≤1 hour post-invasion) ring-stage parasites, another fluorescently tagged version of *Pf*START1, called mNeonGreen-HA_3_-PV6, was observed at the tips of the amoeboid arms ^11^. This discrepancy may be explained by age differences between our amoeboid ring-stage NG-*Pf*START1 parasites and the mNeonGreen-HA_3_-PV6 parasites imaged by Fréville *et al*. ^11^. It is likely that the latter parasites were younger, and therefore still retained some of the *Pf*START1 protein that was present in the invading merozoites. As the ring-stage parasites mature, *Pf*START1 likely moves into the PV and may degrade or become too diffuse in the expanding PV space to be detectable, before being re-synthesised in trophozoites.

We attempted to more precisely resolve the location of *Pf*START1 in merozoites by performing expansion microscopy on *Pf*START1-HA schizonts as commercially available NG-specific antibodies did not work using the expansion method. Here, *Pf*START1-HA appeared to localise to small puncta that did not overlap with established rhoptry, dense granule and microneme markers. This agrees with a previous study which reported that, although the *Pf*START1 localisation pattern was observed to be dense granule-like, it did not appear to overlap with the established dense granule marker EXP2 ^8^. Immuno-electron microscopy of invading *P. knowlesi* merozoites indicates dense granules release their contents into finger-like extensions of the PVM soon after invasion ^31^. In *P. falciparum*, less than 12 minutes post-invasion, the dense granule protein RESA appears to have moved from the dense granules to the parasite periphery and is beginning to be exported into the RBC compartment ^32^. As we observed that NG-*Pf*START1 is secreted into the PV shortly after invasion, we speculate that *Pf*START1 is also stored in dense granule-like organelles. It is possible that there are different populations of dense granules in which different cargoes are stored that are released shortly after invasion to help expand the nascent PVM, perhaps even contributing to the formation the finger-like PVM extensions previously observed by immuno-electron microscopy ^31^.

The role of *Pf*START1 in the expansion of the nascent PVM shortly after merozoite invasion is consistent with the observed effect of inhibiting this protein with compound M-833, which renders newly invaded merozoites unable to expand into larger amoeboid ring-stage parasites ^10^. *Pf*START1 appears highly susceptible to inhibition with compounds WEHI-991 and WJM-715, which have EC_50_s <10 nM in standard 72-hour growth assays ^10,14^. Despite the rapid metabolic turnover of WJM-715 in liver microsomes, several hours after administration to mice, the compound remained at a concentration that would be expected to suppress *P. falciparum* growth *in vitro* ^14^. This prompted further evaluation of WJM-715 in the *P. berghei* mouse model of malaria, but the compound failed to suppress growth during 4 days’ treatment ^14^.

The X-ray crystal structure of *Pf*START1 in complex with WEHI-991 provides important insights into the molecular basis of ligand binding and helps explain the reduced susceptibility of M-833-resistant parasites to amino aryl acetamides. WEHI-991 engages in an extensive hydrogen-bonding network involving residues N309 and N330, mediated by a water molecule. Substitution of either residue with lysine would be expected to disrupt these interactions and consequently reduce compound binding. Whether these mutations also affect binding of the endogenous lipid substrate remains unclear. However, the absence of an observable fitness cost associated with these mutations^10^ suggests that lipid binding and transport may be largely preserved. Comparison of the *Pf*START1 structure with a *Pb*START1 homology model further provides a structural rationale for the lower activity of amino aryl acetamides against *P. berghei* relative to *P. falciparum*. In *Pb*START1, residue S297 occupies the equivalent position to N330 in *Pf*START1. Replacement of asparagine with serine at this position is predicted to weaken key ligand interactions, thereby reducing amino aryl acetamide binding affinity and antiparasitic potency.

The *Pf*START1 structure also explains several structure-activity relationship trends previously observed for amino aryl acetamide analogues against *P. falciparum* ^14^. For example, N-methylation of either the aniline nitrogen or the carboxamide nitrogen markedly decreases antiparasitic activity. Both groups participate in the water-mediated hydrogen-bonding network with N309, N330 and Y328, and methylation likely disrupts these critical interactions. The structure also rationalizes the preference for small substituents, such as fluoro or chloro groups, on the amino aryl ring. In contrast, bulkier substituents such as trifluoromethyl or trifluoromethoxy groups are unlikely to be accommodated within the corresponding binding pocket, resulting in reduced affinity for *Pf*START1. Collectively, these structural insights establish a molecular framework for understanding resistance, species-specific activity, and structure-activity relationships within the amino aryl acetamide series. They also provide a foundation for the rational design of next-generation compounds with improved affinity for *Pf*START1 and potentially broader activity against START proteins from multiple *Plasmodium* species.

To assess whether structural divergence in *Pb*START1 reduced its susceptibility to inhibition by WJM-715, thereby potentially explaining the compound’s lack of efficacy in the mouse model, we replaced the START domain of *Pf*START1 with the corresponding domain of *Pb*START1. Compared to wild-type parasites, these chimeric *P. falciparum* parasites were highly resistant to all *Pf*START1 inhibitors tested, indicating the lipid binding pocket of the START domain of *Pb*START1 was different enough for the compounds to no longer bind with high affinity (S9 Fig). M-833, WEHI-991 and WJM-715 inhibited *Pf/Pb*START1 parasite invasion, indicative of a common mechanism of action in wild-type *P. falciparum* and *Pf/Pb*START1 parasites, which suggested the compounds remained on target against the chimeric protein.

In a previous study, following dosage of mice at 50 mg/kg the plasma concentration of WJM-715 peaked at 1.2 µM at the first sampling timepoint, 15 minutes after administration ^14^ – well below the EC_50_ for *Pf/Pb*START1 parasite growth inhibition *in vitro* (∼3 µM) – then rapidly declined, suggesting WJM-715 may not have remained at a high enough concentration for long enough to suppress the growth of *P. berghei* parasites.

For this reason, we replaced the START domain of *Pbstart1* with that of *Pfstart1* and transfected this chimeric gene into *P. berghei* parasites. As the EC_50_ of WJM-715 against *P. falciparum* parasites is <15 nM – well below the plasma concentration of WJM-715 which was maintained in mice for several hours following daily doses ^14^ – we hypothesised that WJM-715 would suppress the parasitaemia in mice infected with chimeric *Pb/Pf*START1 *P. berghei* parasites. Unexpectedly, this did not occur, and the WJM-715-treated parasites grew as well as the DMSO-treated control parasites. While the resistance of *Pf/Pb*START1 parasites *in vitro* had suggested that structural differences between the orthologues contributed to the failure of WJM-715 in mice, the resistance of *Pb/Pf*START1 parasites *in vivo* showed that this explanation alone was insufficient and indicated that pharmacokinetic properties such as poor bioavailability or short half-life were also responsible. Indeed, as the START1 inhibitors have a static effect on the growth of recently invaded merozoites, rather than a cidal effect ^10^, it is possible that newly invaded parasites whose development was arrested by WJM-715 resumed growth once the concentration of the compound had declined a short time after dosing. Furthermore, differences in target essentiality *in vivo* vs. *in vitro* may mean that inhibition of the START1 protein is less inherently deleterious for *P. berghei* growth than it is in cultured *P. falciparum*. Perhaps in a murine *in vivo* environment, more lipids are available to expand the PV to accommodate the ring-stage parasite than in *in vitro P. falciparum* cultures. Collectively, these findings highlight the utility of interspecies target exchange for predicting and/or explaining the failure of promising *P. falciparum* inhibitors in *in vivo* models.

## Materials and Methods

### Parasite culturing

*Plasmodium spp.* parasites were cultured in human O^+^ RBCs (Australian Red Cross LifeBlood) at 4% haematocrit. Cultures were maintained at 37°C under low oxygen conditions (1% O_2_, 4% CO_2_ and 95% N_2_) as per established protocols ^33^. *P. knowlesi* YH1 ^34^ and *P. falciparum* parasites were grown in RPMI-1640 culture medium supplemented with 0.37 mM hypoxanthine, 25 mM NaHCO_3_, 25 mM HEPES, 20 mg/L (*P. knowlesi*) or 31.25 mg/L (*P. falciparum*) gentamicin and 0.5% (w/v) AlbuMAX II.

### NG-PfSTART1 parasites

A 351 bp 5’ homology block of *Pfstart1* (PF3D7_0104200) was amplified from *P. falciparum* 3D7 genomic DNA with Pf_Bgl_1F and START_2R (S2 Table). The 708 bp sequence for NG was amplified using primers NG_F and NG_R (S2 Table). Primers RC.START_F and RC.START_R (S2 Table) were used to amplify the 1068 bp START domain from a recodonised version of *Pfstart1* (Integrated DNA Technologies). The 462 bp sequence for T2A-BSD was amplified with T2A_F and BSD_R (S2 Table) from a synthetic sequence (Integrated DNA Technologies). The 435 bp 3’UTR of *Pfstart1* was amplified from genomic DNA using primers START_5F and START_6R (S2 Table). All the fragments were sequentially joined to each other using overlapping PCRs and Phusion DNA polymerase to produce a final full length PCR product that was inserted into the pJet plasmid (Thermo Scientific). After Sanger sequencing of the gene fusions, 100 µg plasmid DNA was linearised with *XhoI*, combined with the Cas9/guide RNA complex (Integrated DNA Technologies) (S2 Table) and electroporated into *P. falciparum* 3D7 ring-stage parasites as per ^35^. Integration was selected with 2.5 µg/mL blasticidin S and integration into the *Pfstart1* locus was confirmed by PCR (Fig 1B).

### PfPV1-NG parasites

The *Pfpv1* 5’ homology block was amplified from *P. falciparum* gDNA as two overlapping fragments with primers PV1_1F C PV1_2R and PV1_3F C PV1_4R using Phusion DNA polymerase following manufacturer’s protocols (Thermo Scientific). The overlap site corresponds to where the *Pf*PV1 guide RNA is located, where the overlapping primers PV1_2R and PV1_3F contained synonymous Shld mutations to prevent Cas9 re-cutting the modified locus. The two 5’ homology blocks were PCR sewn together with primers PV1_1F and PV1_4R. The 3’ homology block corresponding to the *Pfpv1* 3’UTR was amplified with PV1_5F and PV1_6R. The synthetic HA-mNeonGreen sequence was amplified and then PCR sewn onto the 5’ homology block with primers PV1_1F and HAglmS_R. The 3’ homology block was sewn to the PV1-HA-mNeonGreen fusion with primers PV1_1F C PV1_6R. The full-length repair template was then inserted into the *EcoRV* site of the pBluescript plasmid (Stratagene). For the *Pf*PV1 guide RNA, two complementary primers with overhangs compatible to the two *BbsI* sites in pDC2-cam-Cas9-U6-hDHFR (pCas9) ^36^ were annealed and ligated into pCas9. Both pCas9 and pBluescript donor plasmids (50 µg each) were transfected into uninfected human RBCs and mixed with trophozoite stage *P. falciparum* 3D7 parasites. Parasites were grown on 2.5 nM WR99210 (Jacobus) for seven days to select for integration. Transfected parasites were observed after approximately two weeks and then FACS-sorted using the mNeonGreen fluorescence. Correct integration into the *Pfpv1* locus was confirmed by PCR (S1B Fig).

### Live cell microscopy

4% HCT cultures of *Pf*PV1-NG and NG-*Pf*START1 parasites were treated with Hoechst (1:2000) for 20 minutes at 37°C. Hoechst was then removed by washing the cells in fresh medium containing 10 mM ascorbic acid and cultures were adjusted to 50% HCT by adding 10 µL cell pellet to 10 µL culture medium containing 10 mM ascorbic acid. 5 µL of 50% HCT culture was dispensed onto a glass microscopy slide and covered with a 50 mm glass coverslip. Parasites were imaged immediately using a Zeiss Cell Discoverer 7 LSM900 microscope, pre-warmed to 37°C with 5% CO2, using the Plan-Apochromat 50× water immersion objective. Images were processed using ImageJ.

### Immunoffuorescence assay

*Pf*START1-HA parasites ^10^ were allowed to settle onto 12 mm coverslips pre-treated with Poly-L-Lysine (Sigma). Cells were fixed for 20 minutes at room temperature with 4% paraformaldehyde and 0.0075% glutaraldehyde in PBS. Cells were then washed in PBS and lysed with 0.1% Triton X-100 in PBS containing 0.1 M glycine. Cells were blocked in 3% BSA, 0.01% Triton X-100 in PBS, and probed with anti-HA (1:500; mouse, Sigma) and anti-EXP2 antibodies (1:500; rabbit)^17^. Secondary antibodies (anti-mouse- and anti-rabbit-Alexa Fluor Plus 594 and 488, respectively; 1:2000) were then used. VectaShield Antifade Mounting Medium (with DAPI, H-1200) was added before sealing the coverslips. Parasites were imaged using a Zeiss Cell Axio Observer, and images were processed with ImageJ.

*Ultrastructure expansion microscopy-indirect immunoffuorescence assay* Ultrastructure Expansion Microscopy ^22,37,38^ of Percoll-purified *Pf*START1-HA (B4 clone) schizonts and gel mounting were performed exactly as described by Ling *et al.* ^39^, with the exception that gels were not blocked and probed immediately with antibodies (in 1× PBS). The following primary antibodies were used to probe gels: rat anti-HA 3F10 (Merck Life Science; #12158167001) at 1:50, mouse anti-RAP1 ^23^ at 1:200, rabbit anti-GAP45 ^40^ at 1:500, mouse anti-EXP2 ^41^ at 1:250, rabbit anti-AMA1 ^42^ at 1:100. The following secondary antibodies/dyes were subsequently used to probe gels at 1:1000: SYTOX™ Blue (Thermo Scientific; S11348), goat anti-rat Alexa Fluor™ Plus 488 (Thermo Scientific; A48262), goat anti-mouse Alexa Fluor™ Plus 594 (Thermo Scientific; A-11032), goat anti-rabbit Alexa Fluor™ Plus 647 (Thermo Scientific; A32733). Imaging was performed on a Zeiss laser scanning microscope 980 with Airyscan 2 using the Plan-Apochromat 63×/1.40 Oil DIC objective. The following settings were used to acquire z-stacks with Airyscan 2: full z-stack per Track, optimal sampling (0.148 µm), bidirectional frame scanning, 35 × 35 × 150 nm pixel size, 2 sampling rate, 0.70 µs pixel dwell time, 850 V detector gain and 2% (445 nm), 2-3% (488 nm), 2% (594 mm) and 2-3% (639 nm) laser power. Images were processed using Imaris 10.2.

### Lattice light-sheet method

Highly synchronous NG-*Pf*START1 late-stage schizonts were separated from uninfected RBCs via Percoll gradient centrifugation. Uninfected RBCs were labelled with PKH26 (MINI26, Sigma Aldrich) as follows: RBCs were washed once in complete RPMI medium by centrifugation (3000 rpm/2 min) and resuspended to 0.5% HCT in incomplete RPMI (without NaHCO_3_ and AlbuMAX II) containing 0.5 µM PKH26. Following a 5-minute incubation at 37°C, the labelled RBCs were washed three times in 1 mL Diluent C (MINI26, Sigma Aldrich) (3000 rpm/2 min). Then, the purified schizont pellet was mixed with PKH26-labelled RBCs to a final 0.1% HCT and ∼60% parasitaemia in a 200 µL volume of imaging medium. The suspension was loaded into an 8-well chambered coverslip (µ-slide 8-well high glass bottom; Ibidi) and allowed to settle for 30 minutes. Imaging was performed on a Zeiss Lattice Lightsheet 7. Excitation light was filtered through a quad-notch filter (405/488/561/640 nm). The 488 nm laser was used to excite mNeonGreen (*Pf*START1) and the 561 nm laser for PKH26, illuminating the sample via a 13.3×/0.44 objective with a light sheet length of 30 µm and thickness of 1 µm. Emission signals were collected via a 44.83×/1 detection objective and split by a 565 nm beamsplitter for projection to a dual camera system. Camera 1 was preceded by a 405/488/561/640 multi-band stop filter, while Camera 2 was equipped with a 500–550 nm bandpass filter. Volumes of 1 µL were scanned at 0.4 µm z-intervals with a 2 ms exposure time per frame for 30 minutes. Raw data were deskewed and deconvolved following previously described protocols ^43^. Final image processing and analysis were conducted using Imaris 10.2 software (Oxford Instruments).

### NG-PfSTART1 mean intensity measurements

The mean intensity of NG-*Pf*START1 post-invasion in the lattice light-sheet dataset was measured by spot detection in Imaris 11.0.1 software. Specifically, tight regions of interest were drawn around the merozoite, with spot sizes of NG-*Pf*START1 defined between 0.8-1.5 µm (approximately the size of a merozoite), and mean intensity measurements of NG-*Pf*START1 were obtained from the defined spots throughout the lattice light-sheet video. The raw mean intensity measurements were normalised by min-max normalisation between values of 0-1 for each invasion event, and all (35) values were combined to generate a mean intensity curve. Raw data is shown in S3 Table.

### P. falciparum construct and protein expression

All gene sequences used were retrieved from the VEuPathDB (accessed through www.plasmodb.org)^44^ from reference strains (3D7 for *P. falciparum*). All genes were synthesized by Genscript (Singapore) unless otherwise stated.

Protein preparation for crystallisation was performed as follows. *Pf*START1, comprising residues 149-394, was engineered to remove five potential N-linked glycan sites by mutation: Asn116, Asn318, Asn319, Asn380 and Asn387 to Gln. Additionally, a C-terminal TEV site followed by a 6× His tag was introduced and the gene synthesized and subcloned into pAcGP67. The construct was then expressed in Sf21 cells and purified from the supernatant via Ni-NTA agarose resin (Qiagen) chromatography, followed by size exclusion chromatography (S75 Increase 10/300 GL, Cytiva) pre-equilibrated with crystallisation buffer (10 mM Tris-HCl, 150 mM NaCl, pH 8.0).

### Protein crystallisation

Purified *Pf*START protein was co-complexed with WEHI-991 at a 1:2 protein:ligand molar ratio. The complex was then dilution to 6 mg/ml in crystallisation buffer, mixed 1:1 with mother liquor and set up in hanging drop crystallisation experiments. *Pf*START1-WEHI-991 crystals grew in 0.2 M MgCl_2_, 0.1 M Tris-HCl, 31% PEG 4K, pH 7.92 and were cryoprotected in 15% glycerol. Diffraction data were collected with the MX3 beamline at the Australian Synchrotron (Clayton, Australia) at 100 K (λ = 0.95372 Å) (PDB ID: 44ZY / PDB_000044ZY). Statistics are in Supplementary Table S4.

### Structure determination and model building

The diffraction data were processed and reduced with the XDS package (Kabsch, 2010) before being scaled and merged using Aimless ^45^ in the CCP4 suite ^46^. Two molecules were estimated in the asymmetric unit using the program Matthews ^47^. The AlphaFold3 ^48^ model of the truncated *Pf*START1 protein was used for molecular replacement using Phaser ^49^. Alternative rounds of structure building and refinement were carried out using Phenix ^50^ and assessed and modified using Coot ^51^. A restraint dictionary was generated for WEHI-991 using Grade2 ^52^ and two copies were manually added into the density after the first round of refinement. A composite omit map was generated using the Phenix package “composite omit map”. Refinement and model statistics are found in Table S4.

### Structure prediction

Prediction of START proteins from *Plasmodium* spp. was performed using the AlphaFold3 server ^48^.

### Pf/PbSTART1 parasites

Primers Pf_Bgl_1F and Pf_2R (S2 Table) specific for *Pfstart1* (PF3D7_0104200) were used to amplify the 444 bp 5’ homology block from *P. falciparum* gDNA using Phusion DNA polymerase following manufacturer’s protocols (Thermo Scientific)(S6 Fig). From *P. berghei* ANKA gDNA, primers Pb_1F and Pb_Spe_2R were used to amplify a 975 bp fragment containing the START domain to the end of the coding sequence of *Pbstart1* (PBANKA_0208900). The *P. falciparum* 5’ homology block and the *P. berghei* START1 domains were joined by overlapping PCR and inserted into the p1.2 plasmid via *BglII* and *SpeI* sites ^53^. The 390 bp 3’ homology block was amplified from *P. falciparum* gDNA in two overlapping fragments with primers START_EcoR1.2 and StART1_BglRepR used for fragment 1 and primers StART1_BglRepF and STLP_KasR used for fragment 2 (S2 Table). The overlapping primers StART1_BglRepR and StART1_BglRepF contained a single base mutation to inactivate an internal *BglII* site and the two fragments were joined together by PCR with START_EcoR1.2 and STLP_KasR. The 3’ homology block was inserted into the p1.2 plasmid containing the *P. falciparum* 5’ homology block and the *P. berghei* START1 domains via *EcoRI* and *KasI* sites to produce the donor plasmid p1.2_*Pf/Pb*START1. This plasmid (100 µg) was linearised with *BglII*, combined with the Cas9/guide RNA complex (Integrated DNA Technologies) (S2 Table) and transfected into *P. falciparum* 3D7 ring-stage parasites as per ^35^. Modified parasites were selected using 2.5 nM WR99210.

### Pf/PbSTART1 knockdown

Ring-stage 3D7 and *Pf/Pb*START1-glmS parasite cultures were obtained by treatment with 5% D-sorbitol ^54^ and adjusted to 0.3% parasitaemia. In a 96-well round-bottom plate (100 µL culture per well; 2% haematocrit), parasites were exposed to 0, 0.25, 1 or 2.5 mM glucosamine in technical triplicate and incubated at 37°C for 72 hours (1.5 intraerythrocytic growth cycles). The resultant trophozoite-stage parasites were then lysed by freeze/thaw and parasite growth was quantified by measuring the activity of parasite lactate dehydrogenase (LDH) released upon cell lysis ^55^. Briefly, 75 µL Malstat reagent (0.166 mg/mL nitroblue tetrazolium, 8.33 µg/mL phenazine ethosulfate, 83.3 mM Tris, 185 mM lactic acid, 0.167% (v/v) Triton X-100, 0.423 mM acetylpyridine adenine dinucleotide) was dispensed into each well of a 96-well flat-bottom plate. 30 µL trophozoite lysate was added to each well and the plate was incubated in the dark at room temperature for 30-60 minutes. The absorbance at 650 nm (OD650) was then measured using a spectrophotometer (Thermo Scientific Multiskan GO). After subtracting the background OD650 of 2% HCT uninfected RBC lysate samples, the OD650 of glucosamine-treated parasite samples was normalised to that of untreated samples (representing 100% growth).

### Growth inhibition assays with 3D7 and Pf/PbSTART1 parasites

In 96-well round-bottom plates, cultures of sorbitol-synchronised ^54^ ring-stage 3D7 and *Pf/Pb*START1-glmS parasites at 0.3% parasitaemia and 2% haematocrit were treated with a 9-point, 2-fold dilution series of WEHI-991, WJM-715, M-833, **13i**, **13h** or **10y** (100 µL culture per well). Additional *Pf*START1-targeted analogues of WEHI-991 were each tested in a 3- or 4-point, 2-fold dilution series, with a maximum concentration corresponding to ∼2x the 72-hour growth EC_50_ value in 3D7 parasites ^14^. After 72 hours’ incubation at 37°C, the resultant trophozoite-stage parasites were lysed by freeze/thaw and parasite growth was quantified by measuring the activity of parasite LDH released upon cell lysis as described above. OD650 values of compound-treated samples were normalised to the DMSO vehicle control (representing 100% growth) after subtracting the OD650 of 2% haematocrit uninfected RBC samples. EC_50_ values were derived from curves fitted in GraphPad Prism (version 10.1.1) using the non-linear regression model “log(inhibitor) vs. response – variable slope (four parameters)”. The mean EC_50_ value was calculated from three biological replicates, each conducted in technical triplicate or duplicate.

### Egress and invasion assay

Egress and invasion assays were performed as described by Dans *et al*. ^9^ using 3D7-Hyp1-Nluc parasites (wild-type 3D7 parasites transfected with a bioluminescent nanoluciferase (Nluc) enzyme which is exported into the RBC compartment ^27^) and *Pf/Pb*START1-glmS parasites. Late-stage schizonts (>40 hours post-invasion) were purified by Percoll gradient centrifugation and treated with a 3-fold dilution series of WEHI-991, WJM-715 or M-833. 30 nM ML10 (egress inhibitor), 100 µg/mL heparin (invasion inhibitor), 3 nM atovaquone (negative control) and DMSO (drug solvent) were included as controls. Assays were conducted in 96-well round-bottom plates, with 100 µL culture at 2% parasitaemia and 1% haematocrit per well. Parasites were incubated at 37°C for 4 hours, during which time egress and invasion occurred. For assays conducted with 3D7-Hyp1-Nluc parasites, cell-free supernatant samples, containing Nluc released during merozoite egress, were then collected. Remaining unruptured schizonts were then lysed by sorbitol treatment and compounds and sorbitol were removed by washing three times in culture medium. Plates were incubated for a further 24 hours to allow the parasites to develop to the trophozoite stage. Trophozoites and RBCs were then lysed by freeze/thaw to release intracellular LDH and Nluc. A background control, consisting of Percoll-purified schizonts that were refrigerated at 4°C during the 4-hour drug-treatment window, was included as described previously ^9^. For assays conducted with 3D7-Hyp1-Nluc parasites, egress and invasion were quantified, respectively, by adding 5 µL cell-free supernatant (collected after the 4-hour treatment window) or 5 µL resuspended trophozoite lysate (collected after freeze/thaw) to 45 µL incomplete RPMI medium (without AlbuMAX II and NaHCO_3_) containing 1:1000 Nano-Glo substrate. The bioluminescence was then measured in relative light units (RLU) using a CLARIOstar plate reader (BMG Labtech). For assays conducted with *Pf/Pb*START1-glmS parasites, the success of invasion during the 4-hour treatment window was measured by quantifying the LDH activity of trophozoite lysates as described above. OD650 or RLU values of compound-treated samples were normalised to the DMSO vehicle control (representing 100% growth) after subtracting the OD650 or RLU of background control samples. EC_50_ values were derived from curves fitted in GraphPad Prism (version 10.1.1) using the non-linear regression model “log(inhibitor) vs. response – variable slope (four parameters)”.

### Drug inhibition assays and SYBR DNA staining in Plasmodium knowlesi

In 96-well U-bottom plates, sorbitol-treated ring-stage *P. knowlesi* parasites at 2% parasitaemia and 1% haematocrit were treated with serially diluted compounds of interest, with a final volume of 100 μL per well. DMSO and 100 nM DHA were included as the vehicle control and positive control, respectively. Parasite growth was measured 48 hours later in the next growth cycle at the late trophozoite/schizont stages by SYBR DNA staining. Following the incubation period, supernatants were removed from each well and well contents were resuspended in PBS. An equal volume of SYBR Safe Stain (0.02% (v/v) in SYBR Lysis buffer (pH 7.5, 20 mM Tris, 5 mM EDTA, 0.008% (w/v) saponin, 0.08% (v/v) Triton X-100)) was added to each well and mixed. After incubating for 45 minutes, plates were read on a fluorometer (BMG LabTech PHERAstar FS) (excitation, 485; emission, 520). The background (uninfected RBCs) was subtracted from all samples, and drug treatments were normalised against untreated parasites to calculate the percent survival of drug-treated parasites. EC_50_ values were derived from curves fitted in GraphPad Prism (version 10.6) using the non-linear regression model “log(inhibitor) vs. response – variable slope (four parameters)”. The mean EC_50_ value was calculated from three biological replicates, each conducted in technical duplicate.

### Pb/PfSTART1 parasites

To make the 5’ homology block specific for the *Pbstart1* locus, a 323 bp sequence encoding the 5’ end of the gene including its 5’UTR was amplified from *P. berghei* ANKA template gDNA with primers Pb_1.3F and Pb_2R (S2 Table, S8 Fig) using Phusion DNA polymerase (Thermo Scientific). The *P. falciparum* START domain and its C-terminal coding region were amplified from *P. falciparum* gDNA using primers Pb-Pf_3F and Pf_SacII4R (S2 Table). These PCR products were joined by overlapping PCR to create a gene replacement block which was inserted into the pL0035 plasmid ^56^ via *ApaI* and *SacII* sites (S8 Fig). To make the 3’ homology block, the last 414 bp of the *Pfstart1* coding sequence were amplified from *P. berghei* gDNA using primers Pb_KpnI5F and Kpn_EcoRV6R and were inserted into the pL0035 plasmid via *KpnI* and *EcoRV* sites. This completed the p35_*Pb/Pf*START1 donor plasmid (S8 Fig). To target this plasmid to the *Pbstart1* locus, 2.5 μg of the plasmid was linearised with *HindIII* and *EcoRI* and then added to 2.5 μg pABR099-1 ^57^, which encodes Cas9 and a guide RNA specific for this locus (S8 Fig, S2 Table). The DNA mixture was then added to 100 μL Amaxa nucleofector solution (Lonza) and electroporated into purified schizonts using a Nucleofector II (Amaxa biosystems) device and program U33 as previously described ^58^. Schizonts were transferred to a 1.5 mL tube (Eppendorf) and 50 μL RPMI medium was added, before samples were drawn up into a 1 mL insulin (29-gauge) syringe and injected intravenously (IV) into 6-week-old female Swiss mice. Transfectants were selected by administering 70 μg/mL pyrimethamine (pH 4.5) to the drinking water of mice 24 hours after transfection. Mice were humanely killed by slow fill CO_2_ and blood was harvested when the parasitaemia of infected mice reached >5%, and genomic DNA was extracted to assess for integration. The *Pb/Pf*START chimericparasite line was cloned out by limiting dilution, where Giemsa stains were used to calculate parasitaemia, and the parasitaemia and HCT were used to calculate infected RBCs/mL. Subsequently, dilutions were made using PBS so that each mouse was injected IV with one parasite using a 29-gauge insulin syringe. Parasites were monitored by tail blood smears at various time points and blood was harvested via cardiac puncture for genomic DNA as previously described ^59^.

### P. berghei parasite lines and propagation

Stabilites of RBCs infected with the *Plasmodium berghei* ANKA strain clone 15cy1 stored in liquid nitrogen were thawed and injected interperitoneally into a donor mouse. The parasite load in animals was monitored by visualising methanol-fixed blood smears stained with 10% Giemsa by microscopy using 100 × magnification under oil. At the desired parasitaemia mice were humanely euthanised by slow fill CO_2_ and blood was harvested.

### Quantitative reverse-transcription polymerase chain reaction (qRT-PCR)

Gene expression was examined in 3D7 (wild-type) or *Pf/Pb*START1 hybrid parasites. Briefly, RBCs infected with schizont-stage parasites were harvested by lysing with 0.15% saponin in PBS. Following RNA extraction using the RNeasy Mini Kit (Qiagen), cDNA was synthesised using QuantiTec cDNA synthesis kit (Qiagen) and then subjected to qRT-PCR using 2×SensiFASTMix™ SYBR® Low-ROX master mix (Bioline). The following primer sets were used: DO2158/DO2159 for *Pf/pbstart1*, DO1812/DO1813 for *exp2* (PF3D7_1471100), DO1810/DO1811 for the house-keeping gene *fructose-bisphosphate aldolase* (*fbpa*; PF3D7_1444800) (S2 Table). The expression levels of *Pf/pbstart1* and *exp2* were normalised against the *fbpa* housekeeping gene and the fold-change calculated using the 2ΔΔCt method (Rao, Huang et al. 2013) to compare the expression of the hybrid line compared to wild-type.

### Compounds used in mouse studies

WJM-715 was supplied from WEHI ^10^, whilst artesunate was purchased from Sigma. Stock solutions of all compounds (10 ×) were made by dissolving compound in DMSO. Each day, a fresh aliquot of stock compound was diluted in vehicle such that the final concentration was 10% compound, 40% Polyethylene glycol 300, 5% Tween 80 and 45% saline for administration to mice. WJM-715 and artesunate solubilised well.

### In vivo efficacy testing

Biological assessment of the *in vivo* antimalarial efficacy of compounds was assessed using the *P. berghei* rodent malaria 4-day suppressive (Peters) test in 6-week-old female mice. Mice were infected intraperitoneally with 2 × 10^7^ RBCs infected with transgenic *P. berghei* expressing chimeric *Pb/Pf*START1. At 2 hours and days 1, 2 and 3 post-infection, mice were dosed with either 50 mg/kg WJM-715 (n=4) or vehicle control (n=4) by oral gavage. One mouse was also dosed with artesunate (Sigma) using the same regime at 30 mg/kg. Note that these experiments were performed alongside experiments testing other compounds for their efficacy at clearing *Pb*ANKA parasites. The parasitaemias of mice were assessed by visualising Giemsa-stained thin blood smears by microscopy and a minimum of 1000 RBCs were counted. Mice were humanely culled the day following last oral gavage. To calculate percent antimalarial activity the following formula was used:

100 − (mean parasitaemia_treated_ day 4 post-infection / mean parasitaemia_vehicle control_ day 4 post-infection) ×100.

### Ethics statement and mice

Female Swiss mice (6 weeks) were sourced from the Animal Resource Centre (Perth, Australia). Rodents were maintained on a standard rodent diet (chow) and housed under controlled conditions at 21°C with a 12:12 hour light: dark cycle. All experiments were approved by the Deakin University Animal Welfare Committee (Project approval no. G03-2023). The use of donated human blood to culture *Plasmodium falciparum* was approved by Alfred Hospital Ethics Committee Project 166/24.

## Supporting information

Supplemental Tanbl1 S1, S2 and S4 and Figures S1 to S9

Full wwPDB X-ray Structure Validation Report

## Acknowledgements

We thank Betty Kouskousis and Burnet Imaging Facility for invaluable assistance. We also thank Lifeblood Biological Resources Australia for providing the human blood and BEI Resources for supplying ML10. We would like to thank the Bio21-WEHI Crystallisation Facility within Melbourne Protein Characterisation at The Bio21 Molecular Science and Biotechnology Institute, The University of Melbourne. This work was supported by the Victorian Operational Infrastructure Support Program received by the Burnet Institute. This work was funded by the National Health and Medical Research Council of Australia (Ideas Grant to P.R.G. 2001073; Development Grant 2014427 to B.E.S.). B.E.S. is a Corin Centenary Fellow.

