## Supplemental Tanbl1 S1, S2 and S4 and Figures S1 to S9 for "*Plasmodium berghei* is resistant to aryl amino acetamides that inhibit *P. falciparum* growth by targeting the phospholipid transfer protein *Pf*START1"

**Title:**

**S1 Table. WEHI-991 analogues used in this study.**

| Compound Name | Compound Structure |
|---|---|
| MMV006833 | 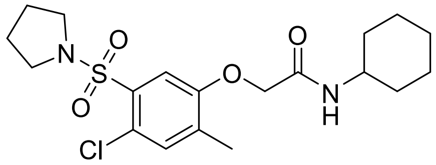 |
| WEHI-991 | 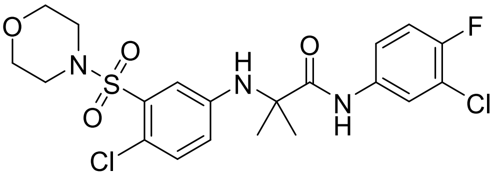 |
| WJM-715 | 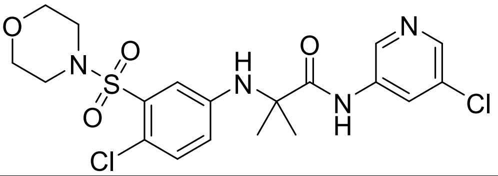 |
| **10r** | 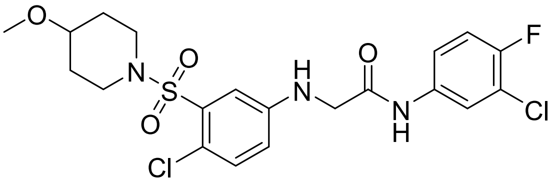 |
| **10n** | 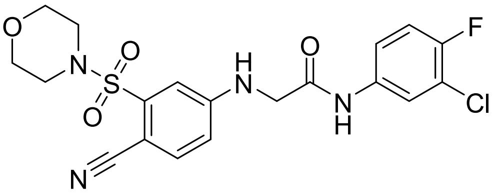 |
| **10q** | 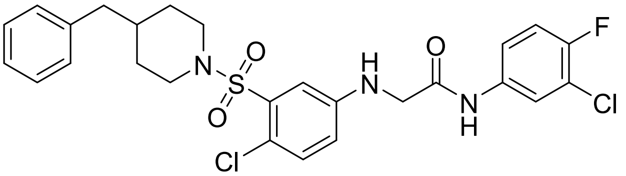 |
| **10k** | 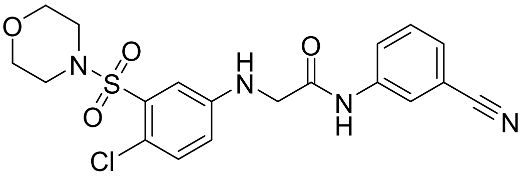 |
| **10b** | 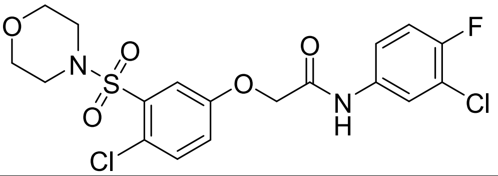 |
| **10f** | 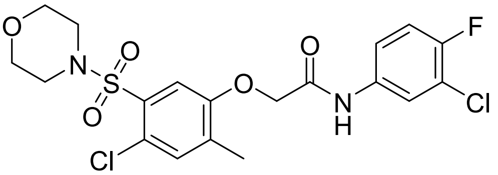 |

| | |
|---|---|
| **13c** | 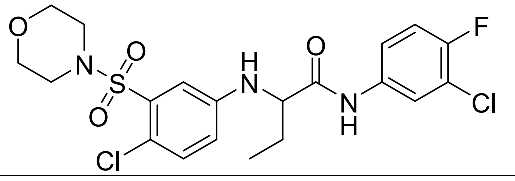 |
| **13j** | 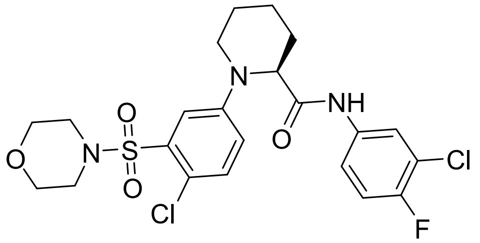 |
| **10y** | 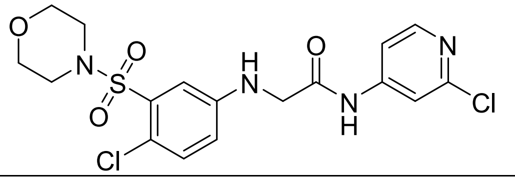 |
| **13i** | 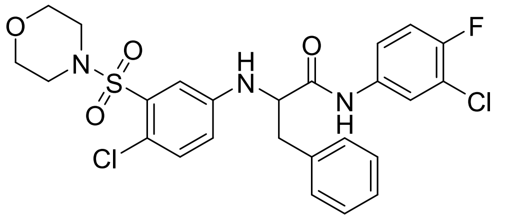 |
| **13h** | 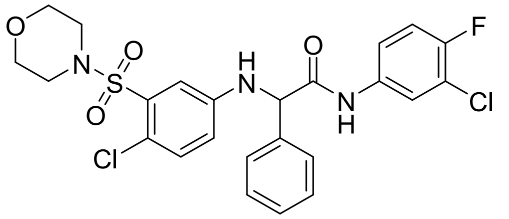 |

**S2 Table. Primers used in this study.**

| Primer Name | Sequence | Purpose |
|---|---|---|
| Pf_Bgl_1F | GGGAGATCTATGAGTTTAAAAAGAAGAAAATTTATTTTGTTGTCTTTTGT | Amplifies 5' *Pfstart1* homology block |
| START_2R | TCCTCGCCCTTGCTCACCATATTATTCCCATCATAAGATTCACCAGT | |
| NG_F | ATGGTGAGCAAGGGCGAGGAGGA | Amplifies mNeongGreen |
| NG_R | AGCCTGCGCTTGGGCCTGAACCTTGTACAGCTCGTCCATGCCCA | |
| RC.START_F | GTTCAGGCCCAAGCGCAGGCT | Amplifies recodonised *Pfstart1* 5' HB |
| RC.START_R | TCCGCTTCCTGATGCactagtGTCTTTGTTAAAAAAAATCCCAAATATC | |
| T2A_F | ACTAGTGCATCAGGAAGCGGAGCT | Amplifies T2A and BSD |
| BSD_R | AGCCCTCCCACACATAACCAGA | |
| START_5F | TTATGTGTGGGAGGGCTAAGAATTCTGCAAATGTGTGATATTGGAAAGGT | Amplifies *Pfstart1* 3' homology block |
| START_6R | GGGGCGCCACACACACATAATACATATATGTATTATTTGGTTCCT | |
| START1_5'_HB_gRNA | AAGCTCAAGCAGAAGCAGAAGGG<br>mA*mA*mG*rCrUrCrArArGrCrArGrArArGrCrArGrArArGrUrUrUrArGrArGrCrUrArGrArArArUrArGrCrArArGrUrUrArArArArUrArArGrGrCrUrArGrUrCrCrGrUrUrArUrCrArArCrUrUrGrArArArArArGrUrGrGrCrArCrCrGrArGrUrCrGrGrUrGrCmU*mU*mU*rU | Guide RNA target site and synthetic sgRNA |
| START_IntF | ATATGCTTTAGGTTATTCGTAGACAGTGT | Used to check for integration into the *Pfstart1* locus |
| NG-Rep-R | AGCAGGGAAACCaGtCCCCTTCACCT | |
| PV1_1F | ACCATTTAACAAGGATAAAGTTTTTTCAACCTTCT | Amplifies first part of *pv1* 5' homology block |
| PV1_2R | CGAATTAATATTCTTAAAAAAGGGAAAAGTGGGTTGGCCA | |
| PV1_3F | CTTTTCCCTTTTTTAAGAATATTAATTCGATCTCTGGA | Amplifies second part of *pv1* 5' homology block |
| PV1_4R | TCGTACGGGTAAGCTGCAGCGCTCGATATTGGTGTGTTTTGATCATTTCCA | |
| PV1_5F | TGAACAAAGAGAAATCACATGATCTTAAATGCCGGACAATTAATAGGACC AATTTGA | Amplifies *pv1* 3' homology block |
| PV1_6R | ACAAAGAACTATACATTCATAGCAAATAATGTGTATTGT | |
| PV1_F_Int | GTAAGTTTTGATGGTCACGATGAACATGT | Used to check integration into *pv1* locus |
| HAglmS_R | AGATCATGTGATTTCTCTTTGTTCA | |
| PV1_sgRNA_F | AAACCTTTCCCATTCTTTAAAAAT | Guide RNAs inserted into BbsI sites of pDC2-cam-Cas9-U6-hDHFR |
| PV1_sgRNA_R | TATTATTTTTAAAGAATGGGAAAG | |
| Pf_Bgl_1F | GGGAGATCTATGAGTTTAAAAAGAAGAAAATTTATTTTGTTGTCTTTTGT | Amplifies the *Pfstart1* 5' homology block from *Pf* gDNA |
| Pf_2R | TAAAGTTGCACTTTTCGTTCTTAAATTTACTGCTGC | |

| | | |
|---|---|---|
| START_EcoR1.2 | GAATTCAGTAAAAATTCATATGATAAAGGAGTAAAATATGTTATGTCCA | Amplifies the *Pfstart1* 3' homology block from *Pf* gDNA |
| STLP_KasR | GGCGCCTTAGTCCTTATTAAAAAATATACCAAATATTTTTTTAAAAAAGTTAACGT | |
| StART1_BglRepF | ACTTAATCAAATATTCCCAGAcCTTATATTTAACTTACA | Used to remove *BglII* site in 3' homology block |
| StART1_BglRepR | TGTAAGTTAAATATAAGgtCTGGGAATATTTGATTAAGT | |
| Pb_1F | AGAACGAAAAGTGCAACTTTAATTACCGACGAATTAGTAACTAAATATTTAAACATTTGT | Amplifies START domain of *Pbstart1* from *Pb* gDNA |
| Pb_Spe_2R | GGGACTAGTTTATTGAAATATACTAATTGTTTTTTGTATAAAACTTGTGCATATTTCA | |
| PfStART_IntF | TGTAATAATTTATACATTTTTATATCAGTTTATTTATTTTTGAGAAGGA | Used to check *P. berghei* START domain has integrated into *Pfstart1* locus |
| HAglmS_R | AGATCATGTGATTTCTCTTTGTTCA | |
| PfSTART-3D7.R | AGAATAAGCTGAAACATCATCCATTTCTCCATATCCT | Combines with Pf-Bgl-1F to check that *Pf* START domain has been replaced in *Pf/Pb*START1 parasites |
| PfSTART1_gRNA | ATGTACAAATTTACATACAT | Sequence recognised by *Pf*START1 guide RNA |
| crRNA sequence | rArU rGrUrA rCrArA rArUrU rUrArC rArUrA rCrArU rGrUrU rUrUrA rGrArG rCrUrA rUrGrC rU | crRNA sequence from IDT |
| Pb_1.3F | CTGACTGGGCCCTCATATAGTAACAGACGATAAATAACGTTAAATTGAAAATCCT | Amplifies 5' homology block from *Pb* gDNA |
| Pb_2R | AAATGCACTAGTTTCATTAATTTTTTTTTTTTCTTCTATTAATTCACT | |
| Pb-Pf_3F | ATTAATGAAACTAGTGCATTTATAAATGATGGTATGTTAGATTATTATTTAAATTTTTGTGA | Amplifies *Pfstart1* domain from *Pf* gDNA |
| Pf_SacII4R | gCCGCGGTTAGTCCTTATTAAAAAATATACCAAATATTTTTTTAAAAAAGTTAACGT | |
| Pb_KpnI5F | GGGTACCACTAATGATATGTATACTCCAGGTTTAGATTATGT | Amplifies 3' homology block from *Pb* gDNA |
| Kpn_EcoRV6R | GGATATCTTATTGAAATATACTAATTGTTTTTTGTATAAAACTTGTGCA | |
| PfSTART_3D7.R | AGAATAAGCTGAAACATCATCCATTTCTCCATATCCT | Combines with Pb_1.3F to show *Pfstart1* domain has replaced *Pbstart1* locus |
| PbSTART_gRNA_R | AAACTAAACCGATACATCATTTAA | The primers combined to encode guide RNA for *Pbstart1* in ABR099-1 plasmid |
| PbStart_gRNA_F | TATTTTAAATGATGTATCGGTTTA | |

| | | |
|---|---|---|
| GU3682 | GATATCATTGAAGCATTTATCAGGG | Sequencing primer for guide RNA insertion into ABR099-1 |
| O2123 | TGTAGCTAAATCGGTTGATCG | Primers used to check correct integration of *Pfstart* domain into *Pbstart1* as per S5 Fig. |
| O2077 | GAAACATCATCCATTTCTCCATATC | |
| O1883 | GATTCATAAATAGTTGGACTTG | |
| O2078 | CATGCGTAATATGGATTGACAATC | |
| O2076 | CATAAAAAGAGTAAGACCGAGAAG | |
| O2123 | TGTAGCTAAATCGGTTGATCG | |
| DO2158 | ACTGGTGAATCTTATGATGGGAATAATG | Amplifies *Pf/pbstart* for qPCR |
| DO2159 | CTAATTCGTCGGTAATTAAAGTTGCAC | |
| DO1812 | GTGGTGGGTATTGTTAGTAAGAG | Amplifies *exp2* for qPCR |
| DO1813 | GTGGCAAAGTTGTTTCTGCATTC | |
| DO1810 | TGTACCACCAGCCTTACCAG | Amplifies *fbpa* for qPCR |
| DO1811 | TTCCTTGCCATGTGTTCAAT | |

**S3 Table.** NG-*Pf*START1 invasion data are shown in a separate Excel file. **Tab 1:** Timing of merozoite invasion steps and secretion time of NG-*Pf*START1 fusion protein. **Tab 2:** The intensity of NG-*Pf*START1 was measured post invasion of RBCs frame by frame by spot detection in Imaris, and normalised (min-max normalisation).

**S4 Table. Data collection and refinement statistics for *Pf*START-WEHI-911.**

| | **PfSTART-WEHI-911** |
|---|---|
| **Beamline** | MX3 |
| **Wavelength (Å)** | 0.95372 |
| **Space group** | $P2_12_12_1$ |
| **Cell dimensions** | |
| ***a,b,c*** **(Å)** | 76.171, 77.012, 86.543 |
| **α, β, γ (°)** | 90.0, 90.0, 90.0 |
| **Resolution (Å)[a]** | 57.53-1.60 (1.63-1.60) |
| **No. molecules in ASU** | 2 |
| **No. observations** | 901,797 (42,088) |
| **No. unique observations** | 67,288 (3,240) |
| **Multiplicity** | 13.4 (13.0) |
| **$R_{merge}$ (%)** | 7.7 (157.2) |
| **$R_{pim}$ (%)** | 2.2 (44.5) |
| **<I/σ I>** | 17.5 (1.8) |
| **$CC_{½}$** | 99.9 (59.9) |
| **Completeness (%)** | 99.3 (97.7) |
| **Refinement Statistics** | |
| **Reflections used in refinement** | 40,445 (2,818) |
| **Reflections used for R-free** | 2,027 (153) |
| **Non-hydrogen atoms** | 4,477 |
| **Macromolecule** | 3,982 |
| **Ligands** | 112 |
| **Solvent** | 383 |
| **$R_{work}$ / $R_{free}$** | 23.35 / 24.45 |
| **Rms deviations from ideality** | |
| **Bond lengths (Å)** | 0.010 |
| **Bond angle (°)** | 1.15 |
| **Ramachandran plot** | |
| **Favoured regions (%)** | 97.66 |
| **Allowed regions (%)** | 2.13 |
| **B-factors (Å$^2$)** | |
| **Wilson B-value** | 24.43 |
| **Average B-factors** | 30.60 |
| **Average macromolecule** | 30.24 |
| **Average ligands** | 23.18 |
| **Average solvent** | 36.49 |

a. Values in parentheses refer to the highest resolution bin.

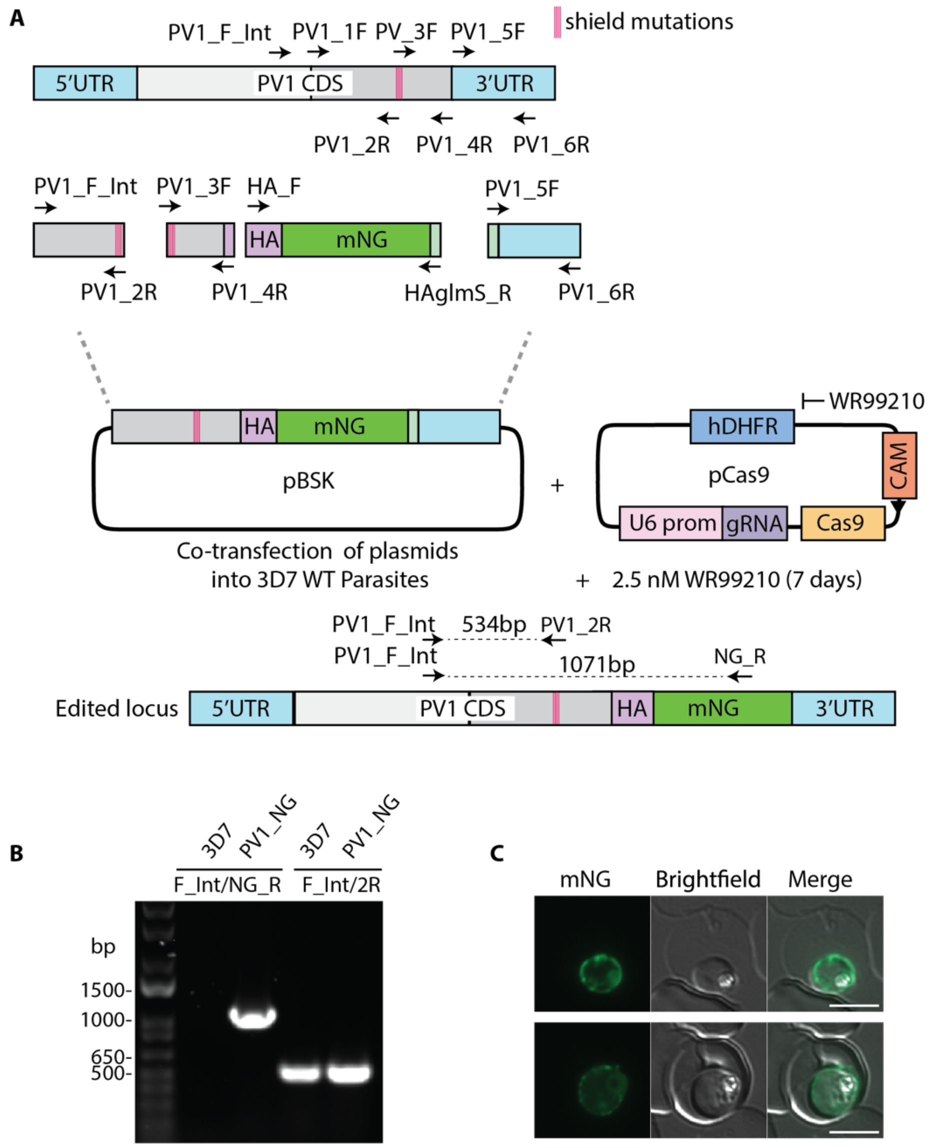

**S1 Fig. Successful tagging of PV1 with HA-mNeonGreen. A)** Diagram showing how *Pf*PV1-HA-mNeonGreen (PV1-HA-mNG) repair template was generated using a multi-step PCR approach and then ligated into a pBSK plasmid. Four shield mutations were introduced via primers (2R and 3F) to prevent Cas9 from cutting the modified gene locus. The repair template was then co-transfected with a second pDC2-cam-Cas9-U6-hDHFR plasmid containing a *Pf*PV1 specific guide RNA into 3D7 WT *P. falciparum* parasites. **B)** Genotyping PCRs confirm successful integration of HA-mNG into the *Pfpv1* locus, where primer PV1_F_Int (F_Int), upstream of the donor block, was combined with primer NG_R to detect insertion into *Pfpv1*. These primers produced no product in parental 3D7 but control primers F_int and PV1_2R (2R) did produce a product in *Pf*PV1_NG and 3D7 parasite lines. **C)** Live cell microscopy confirmed PV localisation of the *Pf*PV1_NG protein. Scale bars = 5 µm.

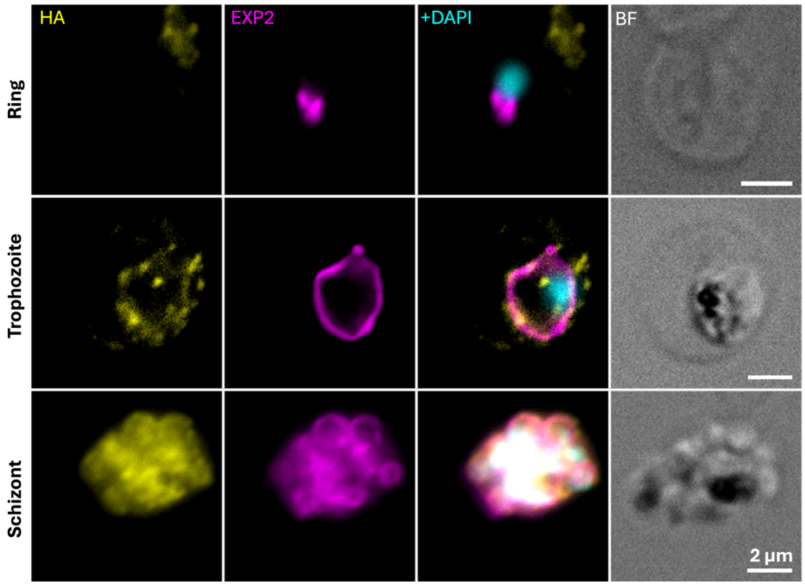

**S2 Fig. *Pf*START1-HA is located within trophozoites and developing merozoites.** Immunofluorescence microscopy was performed on parasites in which the *Pfstart1* gene was appended with a triple haemagglutinin tag as described previously [1]. Fixed parasites of the stages indicated were probed with mouse anti-HA and rabbit anti-EXP2 mAbs and the nuclei were stained with diamidino-2-phenylindole (DAPI). As per the NG-*Pf*START1 parasites, a strong internal signal for *Pf*START1 was detected via the HA tag in schizonts. The signal was weaker in trophozoites and undetectable in ring-stage parasites.

PEXEL
START domain
Drug Resistance mutations: I224F, N309K, N330K

```
PBANKA_0208900.1-p1       MNFKQNKYYIFLVSLLVLDKYN------------------NNAGACKSK-----------    31
PF3D7_0104200.1-p1        MSLKRRKFILLSFVFSVV-ELIFGYD-----NII-----NNYGGLSSVV----------V    39
PKNH_0209300.1-p1         MILKKSKLLALSLLLALV-EYVCRDGRRKWMSLLTSWEVASFDGSNVILNENRRGKYVHI    59
                          * :*: *   : . : :: :                    .  *

PBANKA_0208900.1-p1       --------NSLNEESYIS-----TNL--ISFKNVTNYRILEEALMD----------VENV    66
PF3D7_0104200.1-p1        YNK-NVGINKIGDNVYKN---------------KIKSRILKENKEESLETAAVNENTKDV    83
PKNH_0209300.1-p1         FPRKNASHFARNMTMYRQKERRERNVVCIVPSDISRNRILHEQVEEENESVVREINGNDV    119
                                     .   * .                 . ***.*   :           ::*

PBANKA_0208900.1-p1       KENLKYVQQAKME------ENELNVEGNAQ--------KESPIPSVKSKSELIEEKKKIN    112
PF3D7_0104200.1-p1        KDSCNYSTPSSVFNNDKENTASFIKTGESYDGNNVQAQAQAQAQAEAEGSLENAAVNLRT    143
PKNH_0209300.1-p1         N-----------------------VSGNSDNANDIA-EYEETIKGQASTQEELPQQSVGT    155
                          :                         *::          :    .  . .      .  .

PBANKA_0208900.1-p1       ETSAFITDELVTKYLNICDSVHNNTFIYNNDESFNKSEDCTLHKNLIDEKNNSNYEIIGH    172
PF3D7_0104200.1-p1        KSATLINDGMLDYYLNFCDQVNNKTFPFNECTNLHTSDMVTLCKNNNDEKSKITYDLLGY    203
PKNH_0209300.1-p1         KVSPLIEKDTIDHYLDVCTRIRNKTFSLENCEIFYKSPEVTLYKNILEDKSETRYDLIGY    215
                          : : :* .  :  **:.*  :.*:**  ::   : .*   ** **  ::*.:  *:::*:

PBANKA_0208900.1-p1       GKLNDVSVYGMNYALSNISAIKEWNTHISYLNYLNLTKETIEDKIKRHEKIDPSKYTLND    232
PF3D7_0104200.1-p1        GEMDDVSAYSLSYALNDVETIKDWNKNIYKLNYLNLKKSDIQEKIDKNEKINVNNFILQS    263
PKNH_0209300.1-p1         GTLDDVSLYGGSQALNNIDIIKEWNKNIYKINYLKLNKTSILEKFKDGQKIDPSKYILQN    275
                          * ::*** *. . **.::. **:**.:*  :***:*.*  * :*:.  :**: .:: *:.

PBANKA_0208900.1-p1       AN--MLNNNSYLYLINGLPWPFRSHDAPYEYYQKYFPDKNMILIVNRSIKNAFKDVSGYT    290
PF3D7_0104200.1-p1        EKDVPEDERKYLYLINGLPWPFKSQDTIYEVYQKYYNNKNMLLIINKSLNDVFDNNSSYA    323
PKNH_0209300.1-p1         EH--LRENRRYIYLINGLPWPFKSQDTVYEFYQKYIEDQNILLVANKSVNNVFTDSSYYT    333
                           :    ::. *:**********:*:*: ** ****  ::*::*: *:*:::.* : * *:

PBANKA_0208900.1-p1       RIRNYESFFCLYPKTNDMYTPGLDYVSSILYDVSIPPFLKNSILNQLFPKLILDLNQSSR    350
PF3D7_0104200.1-p1        RINNYENFFCIYPKSKNSYDKGVKYVMSIIYDVNIPKFIQNNILNQIFPDLIFNLHNTSI    383
PKNH_0209300.1-p1         RIRDYENFFCLYPKNKNSYEKGLDYVISVYYDVNIPKFIRNNILGQIFPALIFDLHKSSK    393
                          **.:**.***:***.:: *  *:.** *: ***.** *::*.**.*:** **::*:::*

PBANKA_0208900.1-p1       KYTKVGLSLTDEEKASYLLSLQIKGDNKKLKSVDNKIKNSLNSADNKIKNSLNSADNTEK    410
PF3D7_0104200.1-p1        AITNKTVGTVVDLSKNEQNAWHAHS----LKNVKPE--DTPNTE-----HVQTDPENSYG    432
PKNH_0209300.1-p1         ELTEKGLSLSSEEINKNQLPFQMDN----FSSEDGA--GGLAGM-----DGQVAKSPFFG    442
                            *:  :.   :   .     : ..    :.. .    .          .     .

PBANKA_0208900.1-p1       LNILWIIFVNPVYKIIEICTSFIQKTISIFQ--- 441
PF3D7_0104200.1-p1        LGFIKMIFVDGPYNLWIINVNFFKKIFGIFFNKD 466
PKNH_0209300.1-p1         STVLRVIFVDPFYFIWTTNVNFFKKIYVIVTSIF 476
                            .: :***:  * :    ..*::*   *.
```

PfSTART1 lipid pocket amino acids
L177, L185,
Y198, L200,
L214, L218, I224, W227, N228, N230,
L277, N279, L281, T290, Y292, N309,
Y328, N330, F332,
Y348, M350, I352, Y354,
Q363, L367, I370, F371, L374, I375, L378
Pb or Pk START1 lipid pocket amino acids (above) not conserved with PfSTART1

**S3 Fig. Multiple protein sequence alignment of START1 proteins from *Plasmodium berghei* ANKA, *P. falciparum* 3D7 and *P. knowlesi* YH1.** START domain of *P. falciparum* as defined by [2] is shaded in grey. Amino acid positions that were mutated in *P. falciparum* parasites selected for resistance to compound MMV006833 are highlighted in cyan. Amino acids predicted to line the lipid binding pocket are magenta with differences between *P. falciparum* and the other species in green. Putative PEXEL motifs are highlighted in yellow. Alignment was made in PlasmoDB (https://plasmodb.org/plasmo/app) using Clustal Omega.

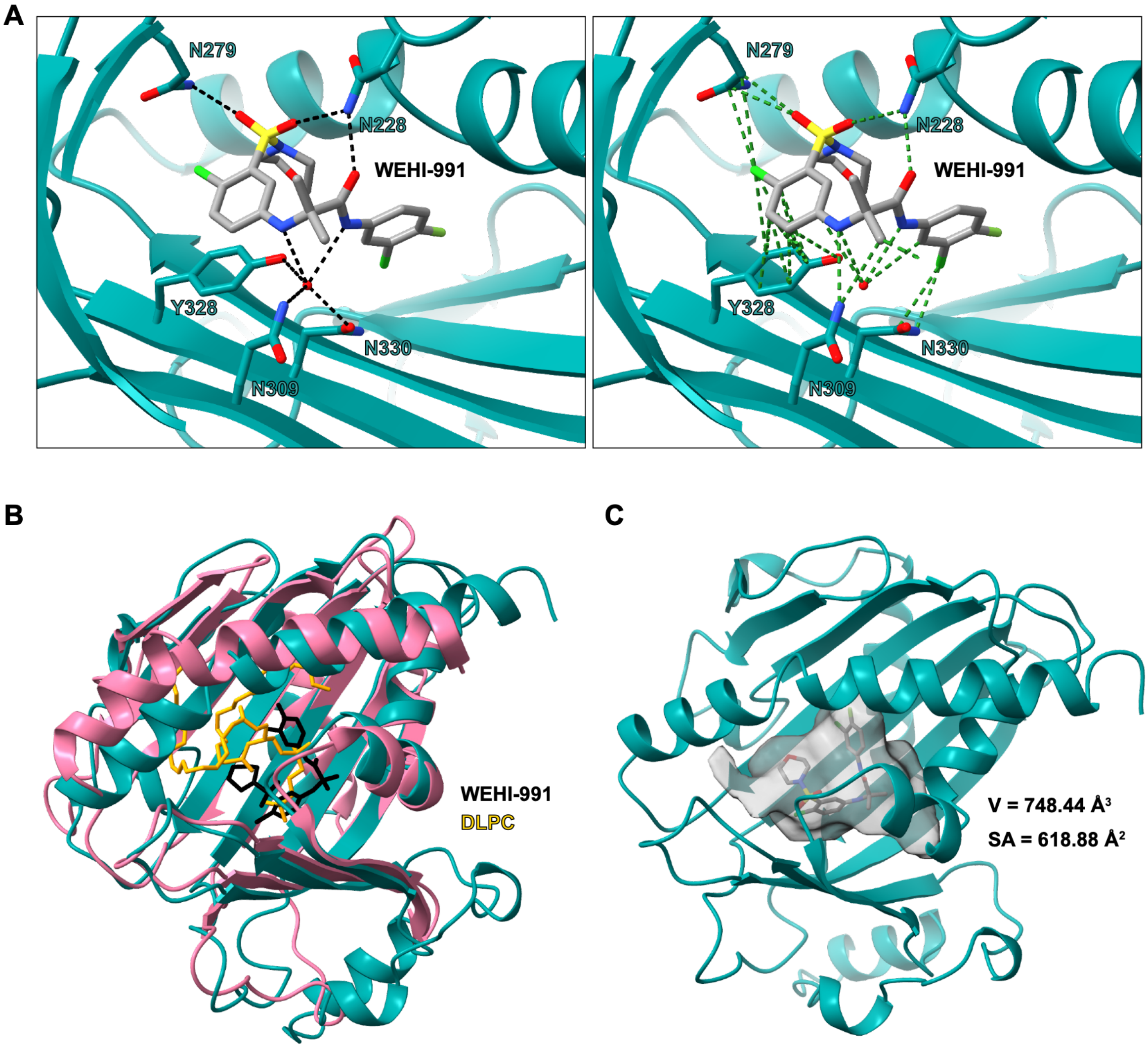

**S4 Fig. WEHI-991 only partially occupies the central lipid binding cavity of *Pf*START1. A)** (left) Hydrogen bonding (black dashed lines) of WEHI-991 to *Pf*START1 and a single co-ordinated water molecule (red circle). (right) All contacts (green dashed lines) of WEHI-991 to *Pf*START1 and a single co-ordinated water molecule (red circle). **B)** The crystal structure of *Pf*START1 (teal) with WEHI-991 (black) overlayed with the crystal structure of human phosphatidylcholine transfer protein (PC-TP). START domain (pink) with 1,2-dilinoleoyl-*sn*-glycerol-3-phosphorylcholine (DLPC, PDB: 1LN2, yellow). **C)** Space-filled central lipid binding cavity of *Pf*START1 (grey) containing WEHI-991. The cavity was calculated to have a total volume of 748.44 Å$^3$ and a surface area of 618.88 Å$^2$.

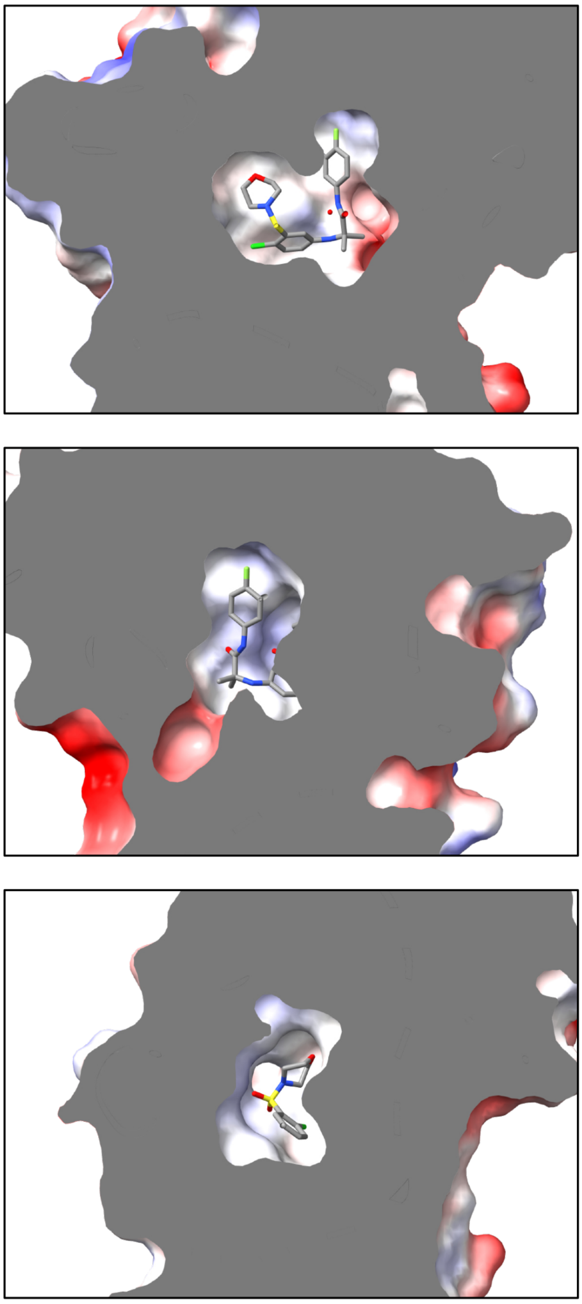

**S5 Fig. The central lipid binding cavity of *Pf*START1 is generally neutral in charge.** Three alternative views of the central lipid binding cavity of *Pf*START1 occupied by WEHI-991. The surface is clipped (grey) to reveal the cavity which is coloured by electrostatic charge (red = negative; white = neutral; blue = positive). Surfaces were generated and coloured in UCSF ChimeraX.

**A**

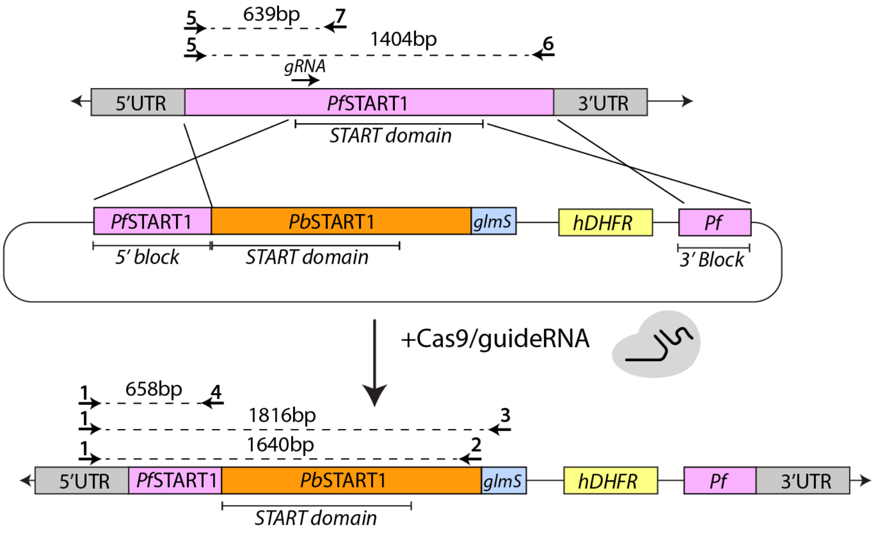

**B**

| Primer pairs | Size (bp) |
|---|---|
| (**1**) PfSTART_IntF and (**2**) Pb_Spe_2R | 1640 |
| (**1**) PfSTART_IntF and (**3**) HAglmS_R | 1816 |
| (**1**)PfSTART_IntF and (**4**) Pf_2R | 658 |
| (**5**) Pf_Bgl_1F and (**6**) STLP_KasR | 1404 |
| (**5**) Pf_Bgl_1F and (**7**) PfSTART-3D7.R | 639 |

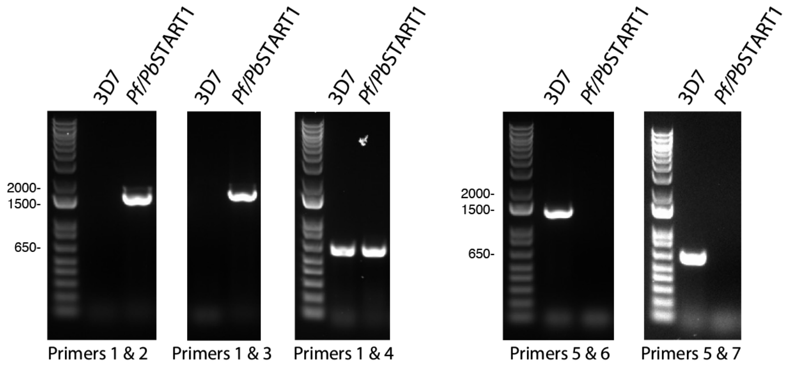

**S6 Fig. The START domain of *Pf*START1 can be replaced with that from *Pb*START1. A)** Diagram of the donor plasmid used to target the *Pfstart1* locus to replace the START domain. Homology blocks derived from the 5' and 3' ends of *Pfstart1* were used to insert the *Pbstart1* START domain into the *Pfstart1* gene. A glmS riboswitch was included in the transfection plasmid to facilitate knockdown of *Pf/Pb*START1. Gene replacement was performed using a Cas9 enzyme and a *Pfstart1* specific guide RNA. **B)** Diagnostic PCRs used to check correct integration into the *Pfstart1* locus using *Pf/Pb*START1-glmS and *P. falciparum* 3D7 parental genomic DNA templates. Primers 1 and 2 and Primers 1 and 3 should only produce a product for the modified *Pf/Pb*START1 locus. Primers 1 and 4 should work with all parasites and Primers 5 and 6 and Primers 5 and 7 should only produce products in the parental 3D7.

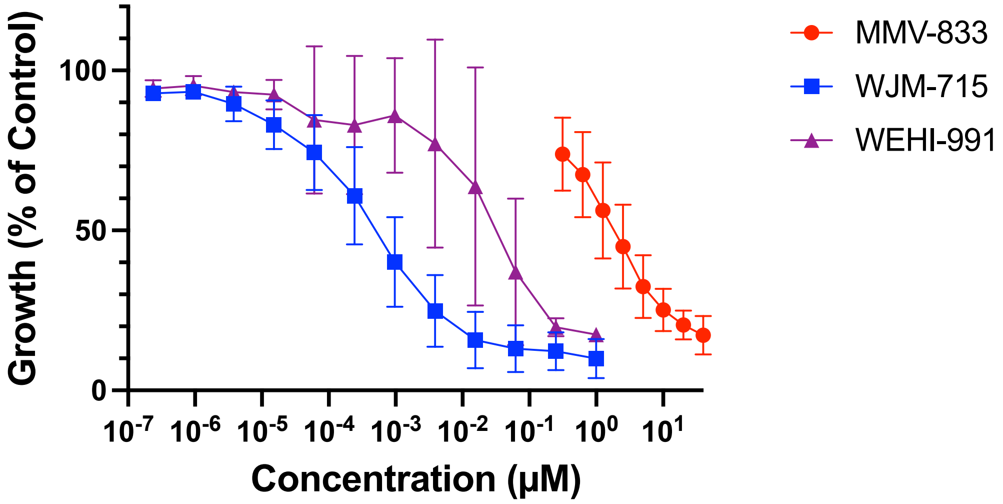

**S7 Fig. *Pf*START1 inhibitors also inhibit the growth of *P. knowlesi* parasites.** *P. knowles*i YH1 parasites were grown for 48 hours in serially diluted *Pf*START1 inhibitors after which their growth was measured through DNA staining with Sybr Green. Growth was measured in three biological replicates (with error bars indicating standard deviation), each conducted in technical duplicate. Curves were fitted and $EC_{50}$s were estimated using Prism V10.

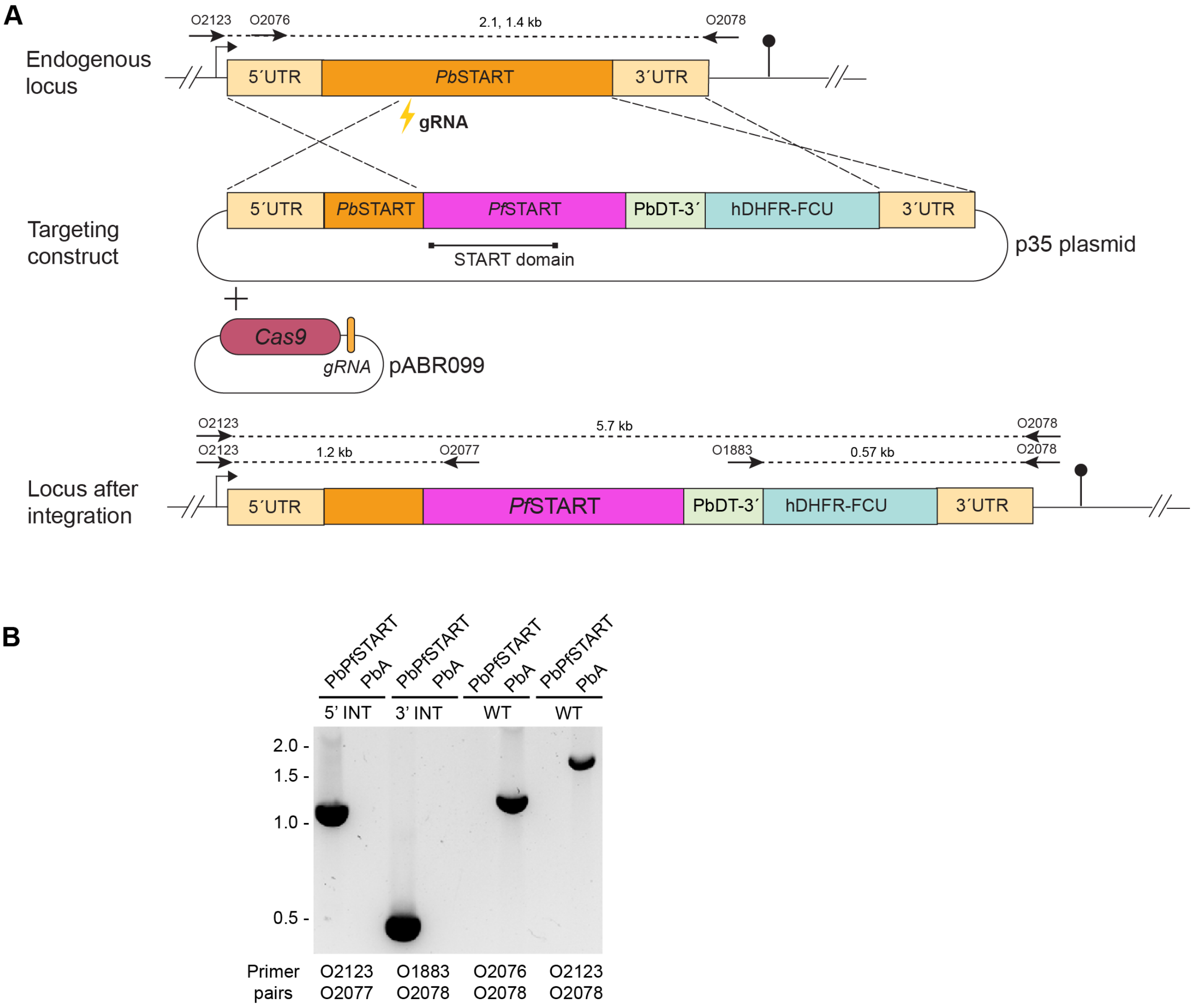

**S8 Fig. The START domain of *Pb*START1 was replaced with the equivalent domain from *Pf*START1 to create *Pb/Pf*START1 parasites. A)** Gene map of donor plasmid used to replace the START domain of *Pb*START1 with that of *Pf*START1 using CRISPR/Cas9. **B)** Diagnostic PCRs used to confirm the gene replacement was successful. The primer binding positions are indicated on the gene maps. PCRs labelled 5'INT and 3'INT should only produce a product in the modified *Pb/Pf*START1 parasites. PCRs labelled WT should only produce a product in WT *P. berghei* ANKA (*Pb*A) parasites.

A

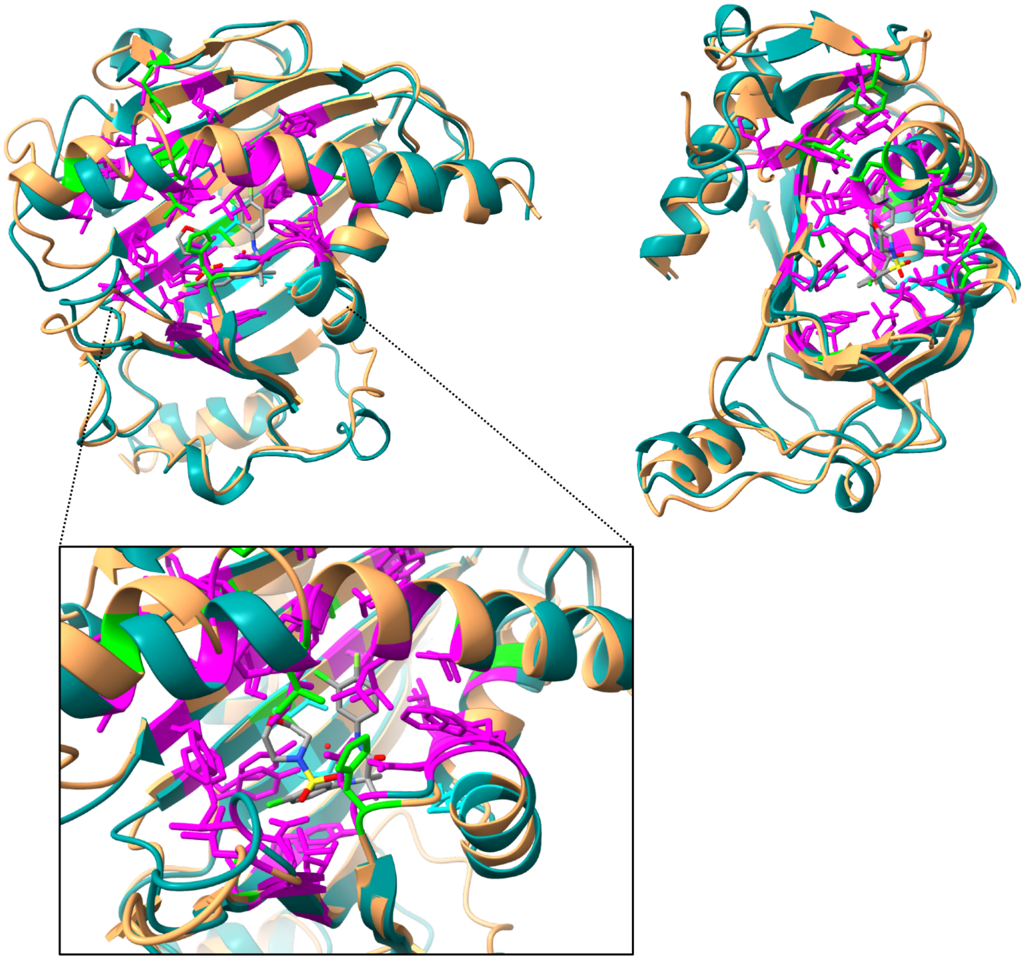

B

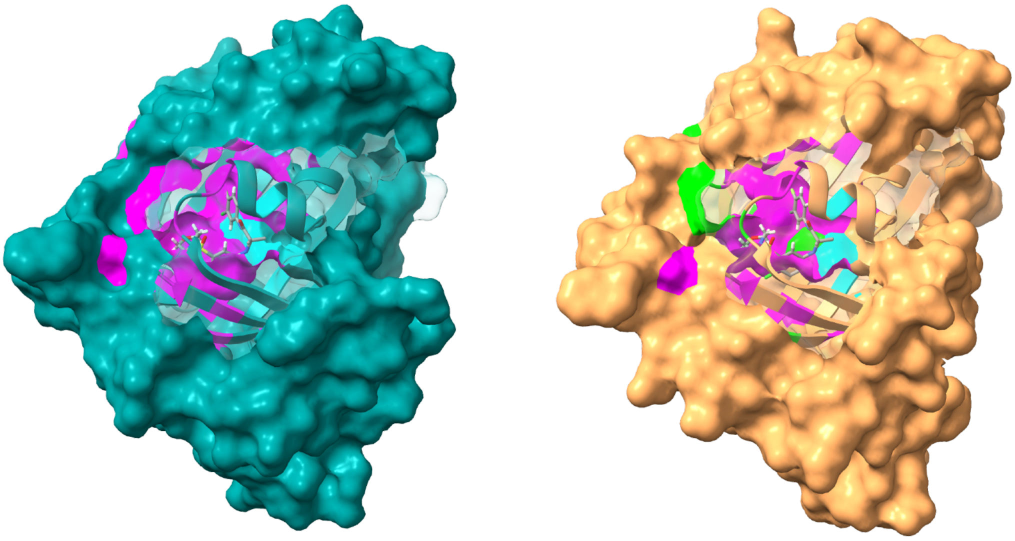

**S9 Fig. Structural visualisation of sequence conservation between multiple species of *Plasmodium* START1 proteins.** Amino acid positions that were mutated in *P. falciparum* parasites selected for resistance to compound M-833 are highlighted in cyan. Amino acids predicted to line the lipid binding pocket are magenta with differences between *P. falciparum* and the other species in light green. **A)** (left) The domain architecture of *Pf*START1 (teal) with WEHI-991 overlayed with the AlphaFold3-predicted structure of the *Pb*START1 START domain (orange). (right) Alternate view of the same structural alignment. Callout: the ligand-binding pocket. **B)** Surface view of *Pf*START1 (teal) and the AlphaFold3-predicted structure of *Pb*START1 START domains (brown). A small volume of surface was made transparent to visualise the central lipid binding pocket.
